# Mechanisms Underlying Economic Revealed Preference Of Multi-Component Bundles In Primate Orbitofrontal Cortex

**DOI:** 10.1101/2021.03.02.433629

**Authors:** Alexandre Pastor-Bernier, Wolfram Schultz

## Abstract

Classic neuro-economic studies suggest that people and animals alike assign values to individual attributes arriving at a total summed valuation and therefore overall appraisal of the offer (Kahnt et al., 2011; Lak et al., 2014). However, further work is needed to determine whether such individual attribute appraisals can predict choice between multi-component bundles. In this paper we show the importance and applications of economic theory to decision neurophysiology concerning multi-component attribute options or bundles. We applied random utility modelling (McFadden, 1973) to choices amongst 10 different two-component bundles in non-human primates (*Macacca mulatta*). We found that behavioural random utility (RUM) complies well with revealed preference theory and recapitulates the fundamental properties of empirically obtained indifference curves (IC). Bundles positioned in the same IC display equal choice preference and similar utility whereas bundles positioned across different ICs display different choice preferences. RUM also captures the characteristic shape of IC on each bundle revealing the synergetic interactions between components. We extended RUM to neurophysiological data and focused our investigation to orbitofrontal cortex (OFC), area 13. Neuronal RUM complied with behavioural RUM at single neuron and population level (N: 54). Neural coding of utility was present at target onset, choice, and reward. However, a distinct and separate group of neurons (N: 26) showed partial compliance with utility in either the chosen or the unchosen bundle options and correlated with both utility and probability of choice. We show that relative chosen-utility coding constitutes a separate type of computation and suggest a role for this type of neuron in value-to-choice processing in OFC.

## INTRODUCTION

Rewards or goods contain typically at least two attributes or components, such as taste and calories. We often trade-in some amount of one component to gain one unit of the other component. Experimentally, these components can be modelled as distinct rewards in bundles of two goods. By trying to obtain the most preferable combination of two goods, we are expressing our preferences (revealed preference) and are aiming to maximize their utility. Our previous studies demonstrate that the revealed preferences and their neuronal correlates in both humans and monkeys reflect the integration of all components into a single scalar variable (Pastor-Bernier et al., 2017; 2020). The OFC in non-human primate and human alike contains neuronal signals for multi-component bundles (Pastor-Bernier et al., 2019; Seak et al., 2021). Revealed preference neurons (RPN) integrated the value from bundle components in the structured manner of revealed preference theory and followed ICs.

Classic economists have long been interested in attribute-based demand models, where it is assumed that consumers derive utility from attributes and characteristics rather than the good s themselves (Lancaster, 1966). This insight serves as the foundation for modeling approaches such as Random Utility (McFadden, 1973). In economics, the class of random utility models (RUM) are routinely applied to capture stochastic choice behaviour within a utility maximization framework (Becker et al. 1963, McFadden 1974, 1981, 2001). Many of these models have the important feature that the probability of choosing an option is related to the difference in utility between alternatives (Mosteller and Nogee 1951, Hey and Orme 1994). RUMs are routinely applied to market supply, including environmental, housing and transportation problems (Adamowicz and Swait, 2011; Carlsson et al., 2012; Costanigro and McCluskey, 2011). Importantly, neuroeconomic work seeking to predict consumer choice (Camerer, 2007; Webb et al., 2013) has led RUMs to physiological type of problems (Smith et al., 2014; Lusk et al., 2016; Webb et al., 2019). However, it remains unknown to what extent RUMs can be empirically validated by the emerging field of neuroeconomics when it concerns choices between multi-attribute options or bundles. Classic neuro-economic studies suggest that people and animals alike assign values to individual attributes arriving at a total summed valuation and therefore overall appraisal of the offer (Kahnt et al., 2011; Lak et al., 2014), but further work is needed to determine whether such individual attribute appraisals can predict choice between bundles. In the present study we seek to understand whether a behavioural risk-less utility representation can predict these choices.

Lusk et al. 2016 is one of the few recent studies to have attempted random utility modeling for bundles of multi-component attributes rather than whole goods. The group conducted fMRI recordings while the subjects were passively viewing bundles consisting of egg cartons, while reported the willingness-to-pay (WTP) for each option presented sequentially. The BOLD signal variation across the subjects accounted for a great deal of the variation in WTP observed, providing evidence that, even in the absence of choice, the brain was evaluating the quality of the items presented. In recent human studies (Pastor-Bernier et al., 2020; Seak et al., 2021) stochastic preferences for bundles containing various amounts of fat and carbohydrates were presented to passively viewing subjects that performed a BDM auction. ICs represented\ revealed stochastic preferences for the bundles presented systematically (Pastor-Bernier et al., 2020) and the BOLD signals in the reward-related brain structures, including striatum, midbrain and medial orbitofrontal cortex followed the characteristic pattern of ICs: similar responses along ICs, but monotonic changes across ICs (Seak et al., 2020).These brain structures integrated multiple reward components into a scalar signal and reflected the synergies between the components, thus corroborating previous observations for bundles consisting of fat and carbohydrates (DiFeliceantonio et al., 2018). Previous studies in non-human primates indicate that the primate OFC encodes revealed preferences of multiple reward options (Pastor-Bernier et al., 2017; 2019; Ferrari-Toniolo et al., 2021) and striatal dopaminergic structures encode economic utility (Stauffer et al., 2016), although it remains unclear whether a representation like riskless utility is explicitly encoded in primate OFC in a multi-component choice scenario. By conducting electrophysiological recordings in area 13 we undertook the task of investigating whether a neuronal correlate for such a representation does exists, while taking account of alternative value representations which could be instrumental in transforming the signal from value to choice.

## RESULTS

### Behavioral comparison between random utility and indifference curves

Random utility modelling on behavioral data was obtained for 10 bundles and 2 monkeys in a total of 32464 trials (average 1910 trials/bundle, Table 1). Behavioral indifference curves (IC) were obtained according to a stepwise procedure (Star Methods, Figure 1 A-D). Three fundamental bundles: blackcurrant vs grape, blackcurrant vs water (Subject 1) and blackcurrant vs mango (Subject 2) were additionally studied with electrophysiological recordings. Titration for random utility was performed in two directions. In each bundle, the blocks in which the titration direction was the vertical axis (Figure 1 G) were used to infer the random utility of the reference component blackcurrant. The blocks in which the titration direction was the horizontal axis (Figure 1 F) were used to infer the utility of the alternative component: grape, IMP-grape, water, apple, lemon, saline, strawberry, mango or peach.

**TABLE 1.** Goodness-of-fit in behavioural RUMs.

|  | **Trials** | **Titration steps** | **AIC** | **Log-Likelihood** | **McFadden R<sup>2</sup>** | **L. ratio test Chisq (p<2.22e-16)** |
| --- | --- | --- | --- | --- | --- | --- |
| **SUBJECT 1** |  |  |  |  |  |  |
| Blackcurrant |  |  | 1851.955 | -918.98 | 0.31688 | 852.59 |
| Blackcurrant -MSG | 3497 | 61 | 753.390 | -369.69 | 0.46662 | 646.85 |
| Water | 1073 | 20 | 4926.107 | -2456.10 | 0.19396 | 1182.00 |
| Grape | 4396 | 61 | 212.993 | -99.496 | 0.47125 | 177.36 |
| IMP-Grape | 2375 | 52 | 4443.631 | -2215.80 | 0.24367 | 1427.8 |
| Strawberry | 1073 | 20 | 275.501 | -130.750 | 0.38886 | 166.39 |
| Apple | 1093 | 32 | 2081.716 | -1034.9 | 0.35825 | 1155.4 |
| Mango | 403 | 12 | 2193.5 | -1090.8 | 0.14959 | 383.74 |
| Lemon | 4236 | 91 | 2180.868 | -1083.40 | 0.16426 | 425.9 |
| Saline | 1067 | 44 | 501.4959 | -243.75 | 0.22942 | 145.14 |
| Blackcurrant † | 271 | 4 | 2845.288 | -1420.60 | 0.3538 | 1555.6 |
| Grape† | 3658 | 49 | 2922.285 | -1459.10 | 0.5082 | 3015.9 |
| Water† | 1806 | 21 | 323.796 | 33.64 | 0.0952 | -159.9 |
| **SUBJECT 2** |  |  |  |  |  |  |
| Blackcurrant | 3421 | 49 | 953.3994 | -469.70 | 0.28342 | 371.55 |
| Peach | 264 | 16 | 225.6232 | -105.81 | 0.34991 | 113.91 |
| Mango | 1881 | 45 | 7698.336 | -3842.20 | 0.21564 | 2112.6 |
| Blackcurrant † | 1700 | 22 | 1264.400 | -629.20 | 0.5186 | 1355.7 |
| Mango† | 250 | 13 | 2133.746 | -1064.90 | 0.4596 | 1811.3 |
| Total | 32464 | 612 |  |  |  |  |

†Goodness-of-fit for utility constrained by bundles containing single components (no interaction)

**Figure 1.**
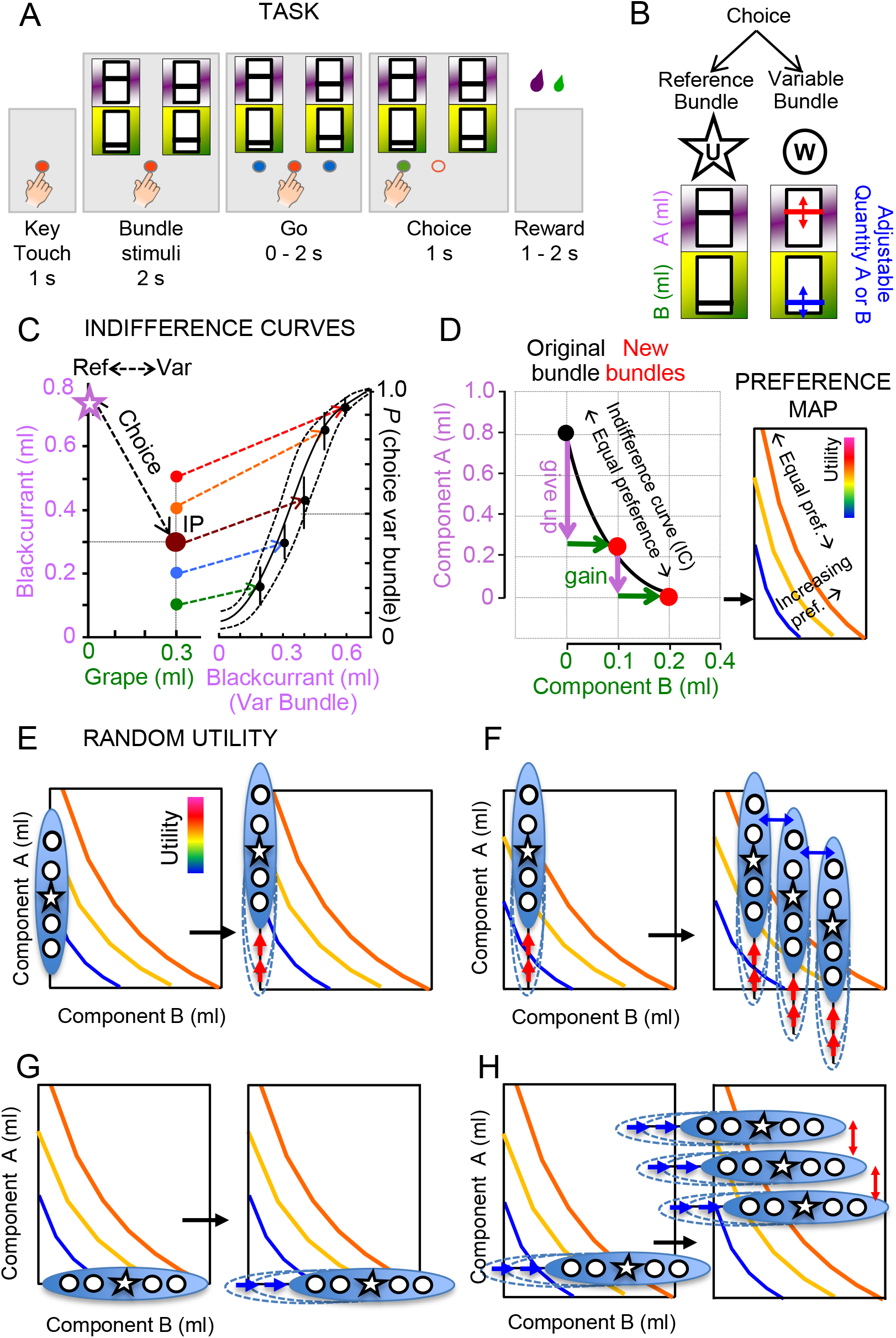
Experimental design and RUM titration process. (A) Behavioral task. After key touch, visual stimuli representing two bundles composed of specific quantities of two liquid goods appear on a computer touch monitor. Their left and right positions relative to the touch key alternate pseudo-randomly. Two lateral round stimuli indicate the signal to GO. The animal releases a contact-sensitive key and touches the stimulus of the bundle of its choice after and then receives the two liquids from that bundle sequentially. Liquid A is delivered first followed by liquid B 500ms after. An interval of 1.6±0.25 s elapses between trials. Each bundle contained the same two liquids with independent quantities (pink, green); the vertical bar position within each rectangle indicates physical quantity. (B) Visual presentation of two-liquid bundles for binary choices. The experimenter sets the quantity of liquids A and B in the reference bundle (‘U’). The quantity of liquid A or B was varied for psychophysical assessment in the variable bundle (‘W’). The reference and variable bundles were visually indistinguishable, apart from their quantity variations. (C-D) Titration procedure for indifference curves. On each given block (C), a point indifference (IP) is obtained by psychophysical titration of the reference bundle against step-wise increasing alternative bundles. (D) The connecting IP define regions of space of equal preference (IC) and increasing preference (maps). (E-H) Titration procedure for random utility. In (E), the RUM dataset for component A (e.g. blackcurrant) consists of a series of binary choices between bundles with a null amount of component B. The dataset is organized in trial blocks (blue ellipse) that present binary options consisting of a reference bundle (star) presented against any of the other 4 available options (circles) positioned at symmetrical equidistance in the Y-axis. The block procedure is repeated iteratively for different reference bundles with gradually increasing magnitude in A (red arrows). The dataset is used by the multinomial logit RUM to obtain the net utility and the random utility estimate of component A with no interactions. (F) Trial blocks containing binary option sets (blue ellipse) where some fixed amount of B is present on a given block. Several blocks with fixed amounts of component B are presented iteratively, mapping the utility space of both components A and B. In this case the multinomial logit RUM yields the net utility and random utility for component A where component B is also a covariate. (G) And (H) represent the RUM dataset for component B without component A interaction, and the RUM dataset for component B with component A interaction, respectively.

In addition to the two sampling directions, we conducted two alternative titration methods to obtain the utility of bundle components with and without interaction.

Blocks containing binary option sets with single-component bundles were used to infer the utility for either component (A or B) without interaction (Figure 1 E and G). Blocks containing binary option sets with non-null quantities in both components were procedurally used to infer the utility of either component (A or B) with interaction (Figure 1 F and H).

Figure 2, S1, S2 show the indifference maps for all bundles tested behaviourally (left column) along with the net utility choice probability (middle column) and the resulting random utility function (right column). In the bundle blackcurrant vs grape (Figure 2A, right column), the utility for blackcurrant juice has a positive slope and is slightly concave, whereas the utility of grape is nearly linear with a much shallower slope (Figure 2B, right column). The variation in the slope of the utility functions depended on both the bundle type and the subjective preferences of the subject. Figure 3 A-D summarizes these results across all bundles and subjects tested.

**Figure 2.**
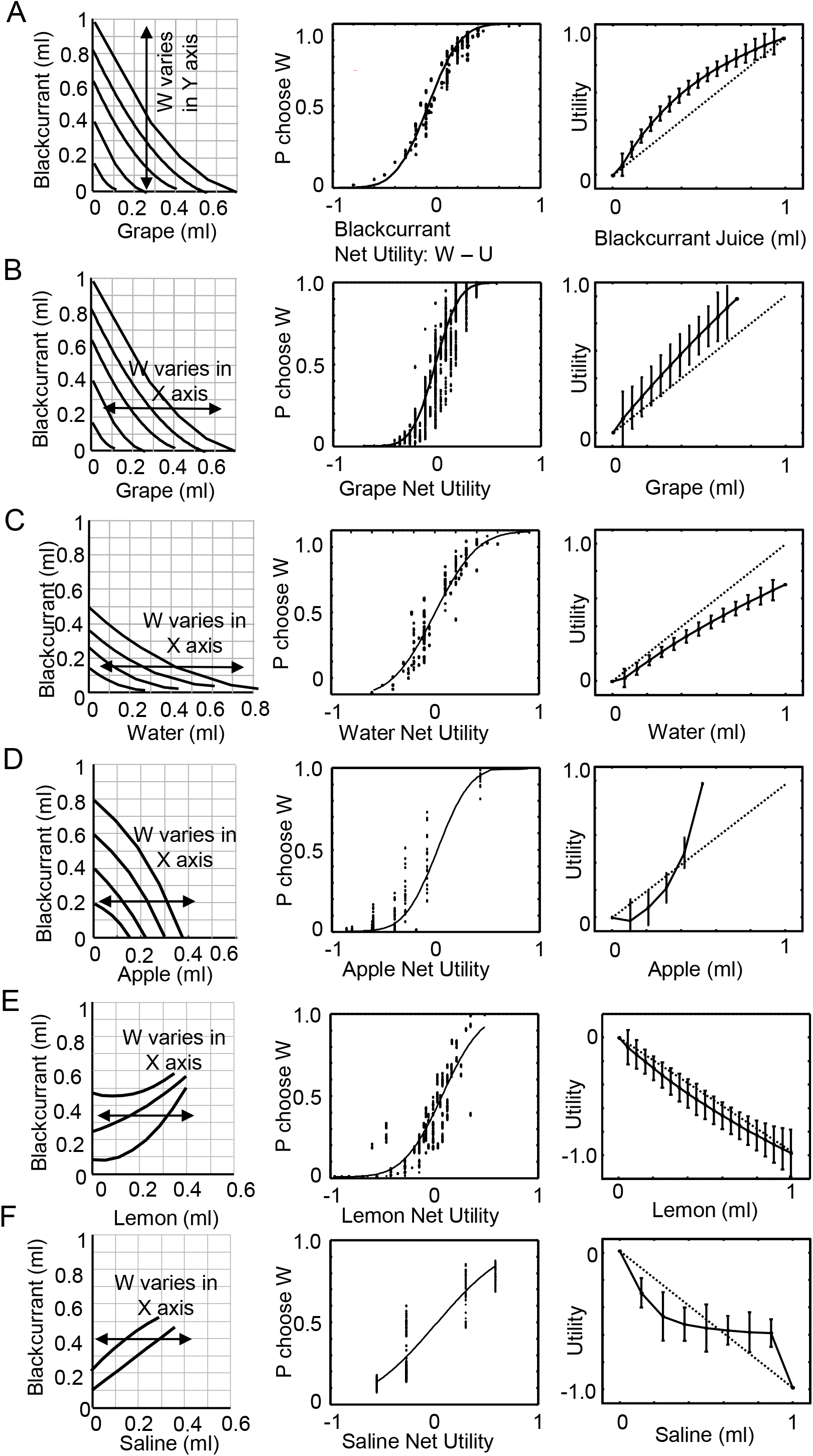
Random utility modeling on behavioral data. (A) Blackcurrant juice vs grape. The left panel shows the indifference map for the bundle blackcurrant vs grape. The double-headed arrow indicates the direction of sampling for inference of utility of Blackcurrant. Blocks of choice trials with bundles with a fixed amount of grape and variable amounts of blackcurrant were used to infer the utility of Blackcurrant. The middle panel shows choice probability as a function of the net utility between the reference bundle (U) and the variable bundle (W). The right panel shows the utility function of blackcurrant with respect to the physical amount of blackcurrant juice (ml). The diagonal dotted line in the right panel serves to show deviation from linear utility defining risk attitude as risk averse (concave) or risk seeking (convex). (B) shows the sampling procedure for the utility of grape, the probability of choice and utility function of grape. (C) refers to the utility of water, (D) apple, (E) lemon, and (F) saline. (E, F) show negative slopes for the utility functions in the bundles blackcurrant vs lemon and blackcurrant vs saline, indicating disutility for consumption of these goods.

**Figure 3.**
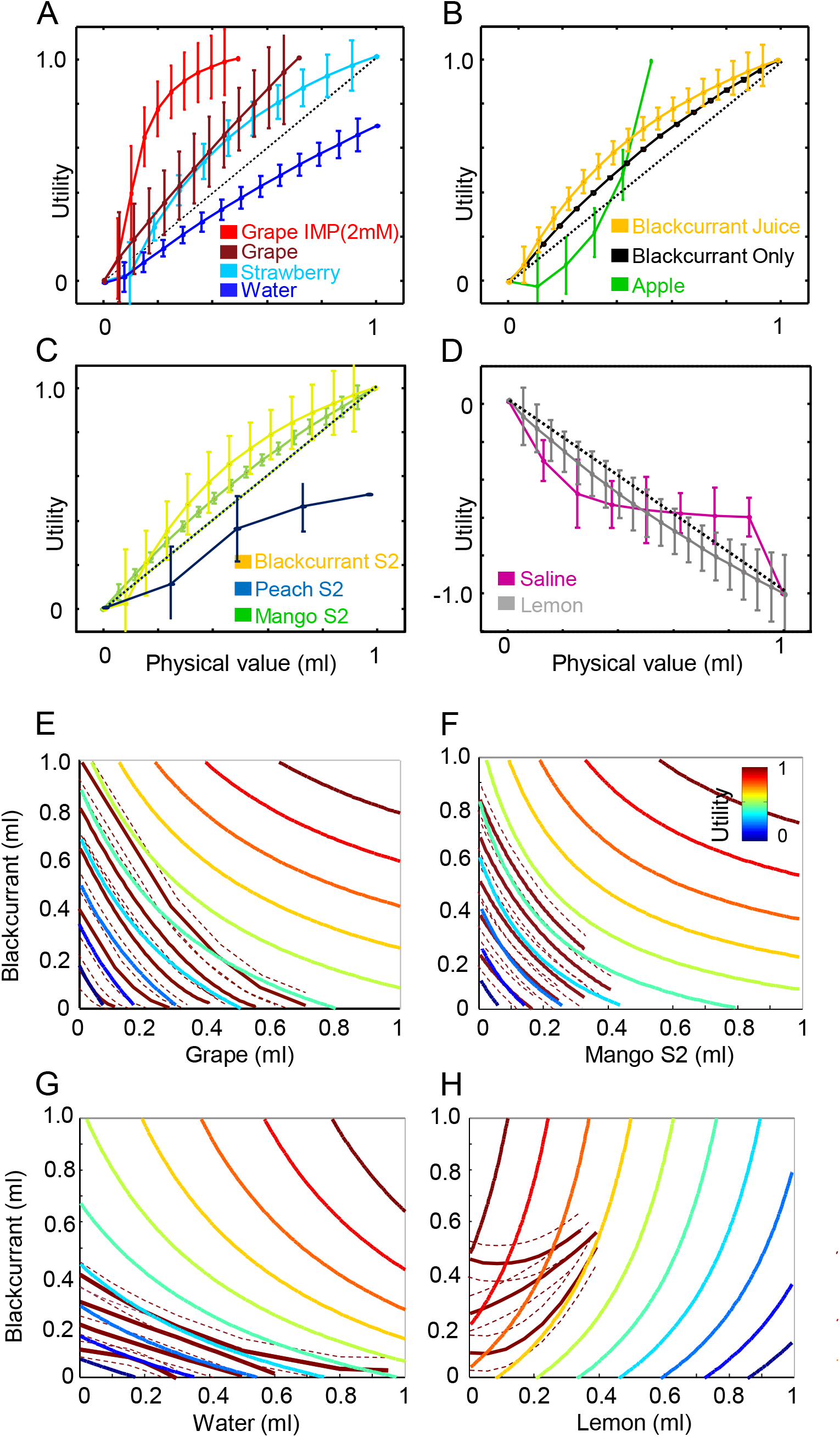
The main features of bundle indifference maps can be explained by the random utility of the components. (A-D) show the utility functions across 12 different bundles (colors) and 2 monkeys. The dotted black line in each panel is shown as reference for a completely linear utility. (A-C) represent positive utility and (D) represent negative utility functions. Random utility errors are shown for each utility estimate with a vertical line. Utility maps along with empirically obtained indifference maps are shown for comparison in (E-H) for 4 representative bundles: blackcurrant vs grape, blackcurrant vs mango, blackcurrant vs water and blackcurrant vs lemon. Empirically derived indifference curves and confidence intervals are shown in Bordeaux with red and dotted lines.

For two bundles, blackcurrant vs lemon and blackcurrant vs saline (Figure 2 E and F, respectively), the utilities had a clearly negative slope. These two components: lemon and saline, introduced a utility cost over the reference component blackcurrant. This result is consistent with the positive slopes of the correspondent indifference curves in figure 2 E-F (left panels). For each increment of lemon or saline consumed an additional quantity of blackcurrant had to be offered to obtain choice indifference. Table 1 summarizes the goodness-of-fit utility calculations for the 10 different bundles tested across two subjects.

We compared random utility estimates and empirically determined indifference maps by combining the utility functions for each separate component in the bundle to reconstruct two-dimensional utility maps for the two-component bundles (Figure 3 E-H). This approach allows for a direct visual comparison between modelled iso-utility lines and indifference curves. As shown previously (Pastor-Bernier et al., 2017), convex indifference curves represent components were the goods acted synergically: for the same quantity of juice, bundles with non-null combinations of the two components have higher value than bundles consisting of single components located in the X or Y axis IC-intercepts. Most bundles tested followed this pattern except for blackcurrant vs apple in Figure 2D. In this case the indifference curves are concave, indicating that bundles with two non-null components are less valuable than their quantity-matched single component counterparts.

Using the modelled utility functions resulted generally in reasonably good fits in comparison with IC features: the currency (IC slope) and synergy (IC convexity) between components. The isolines fell within the confidence intervals of the ICs with exception for the bundle blackcurrant vs lemon (Figure 3 H) which clearly deviates in the two lower bottom IC. This result is consistent with the general measures of goodness of fit for utility with IC empirical data. RUM produced a good fit for most bundles (Table 1), with the best fit obtained for the grape component (McFadden R^2^: 0.47125), followed by mango (R^2^: 0.21564), water (R^2^: 0.19396) with lemon being the weakest (R^2^: 0.16426). Despite random utility and indifference curves being completely different approaches, we observed a statistically significant R^2^ correlation between measures obtained with IC hyperbolic fits and RUM to the empirical data (Rho: 0.5656, p: 0.0350). These results suggest an overall consistency between these two methodologically distinct procedures.

### The shape of the utility function is invariant to the interaction procedure

RUM is very useful for bundle components which are indissociable from each other or introduce a disutility: cost, risk attributes, or non-preferred goods, like lemon or saline. In the later, the utility of either lemon or saline covaries with a non-null amount of a preferred component, like blackcurrant. By doing so the utility of one component in the bundle is inextricably bound to the utility of the other component. However, this design does not let us know how much of the shape of the iso-utility curves can be explained by the interaction of the components or alternatively the intrinsic shape of utility functions for the separate components. Addressing this question is important because it investigates whether the non-linear properties of ICs can be dissociated or predicted from the intrinsically properties of random utility in the components. Answering this question has also bearing on the empirical method used to investigate choice preference in multi-component bundles, because titration of the utility of single components (X or Y axis) is far less laborious than titrating different proportions of both, which is the essence of indifference curve characterization. To address this important question, we estimated random utility in single preferred components that had a good amount of data: blackcurrant, grape, mango and water.

These single-component bundles were used to obtain random utility without interaction in the four control bundles shown in Figure S1A, S2 A-D and Table 1 (controls). We then reconstructed two-dimensional utility maps from single component utility functions (Figure S2 E). The goodness of fit of the random utility models without interaction provided reasonably good fits for the bundles blackcurrant vs mango (McFadden R^2^: 0.4596), blackcurrant vs grape (R: 0.5082) and bundle blackcurrant vs water (R^2^: 0.0952) being the weakest. The fits were also corroborated by AIC and log-likelihood values. These results indicate that in 2/3rds of the bundles tested, random utility of separate components was enough to capture the overall shape features of the indifference curves suggesting that the shape of the utility function could be responsible for the shape of the ICs, and not a consequence of the titration method used, with or without interaction. This is important because it means that the two-component interaction properties that be derive from the shape of ICs in these bundles can be at least partially explained by the intrinsic utility representations of single components.

### Neuronal coding of random utility

We collected 191 neurons in the fundamental bundles: blackcurrant vs grape (81 neurons) and blackcurrant vs water (50 neurons) in Subject 1, and blackcurrant vs Mango (60 neurons) in Subject 2. 105 neurons complied with the characteristics of revealed preference RPN (Star Methods). We define as positive coding neurons those units that integrate both components of the bundle chosen and respond monotonically (more is better) with increasing response for bundles of higher value. Negative coding neurons respond inversely with the value of the option chosen. We recorded 79 positive coding RPNs and 26 negative coding neurons across three bundles. 28 neurons complied with revealed preference (22 positive and 8 negative coding neurons) in the bundle blackcurrant vs grape in Subject 1. Another 35 neurons (25 positive and 10 negative) complied with revealed preference in the bundle blackcurrant vs water, and the remaining 40 neurons (32 positive, 8 negative) complied with revealed preference in the bundle blackcurrant vs mango in Subject 2. Figure 4A-D shows utility functions and response histograms for two single-neuron examples aligned at target onset (left) in the bundle blackcurrant vs grape. The color-coded responses follow monotonically the utility of selected options with increasing physical value (violet lowest utility to red highest utility). In order to compare whether neurons responded best to non-linear random utility function or linear physical value; we regressed the z-scored responses with both variables separately. In the two single neuron examples shown in Figure 4 both regressions of neuron responses to physical value (red dashed line) and random utility (black solid line) were statistically significant.

**Figure 4.**
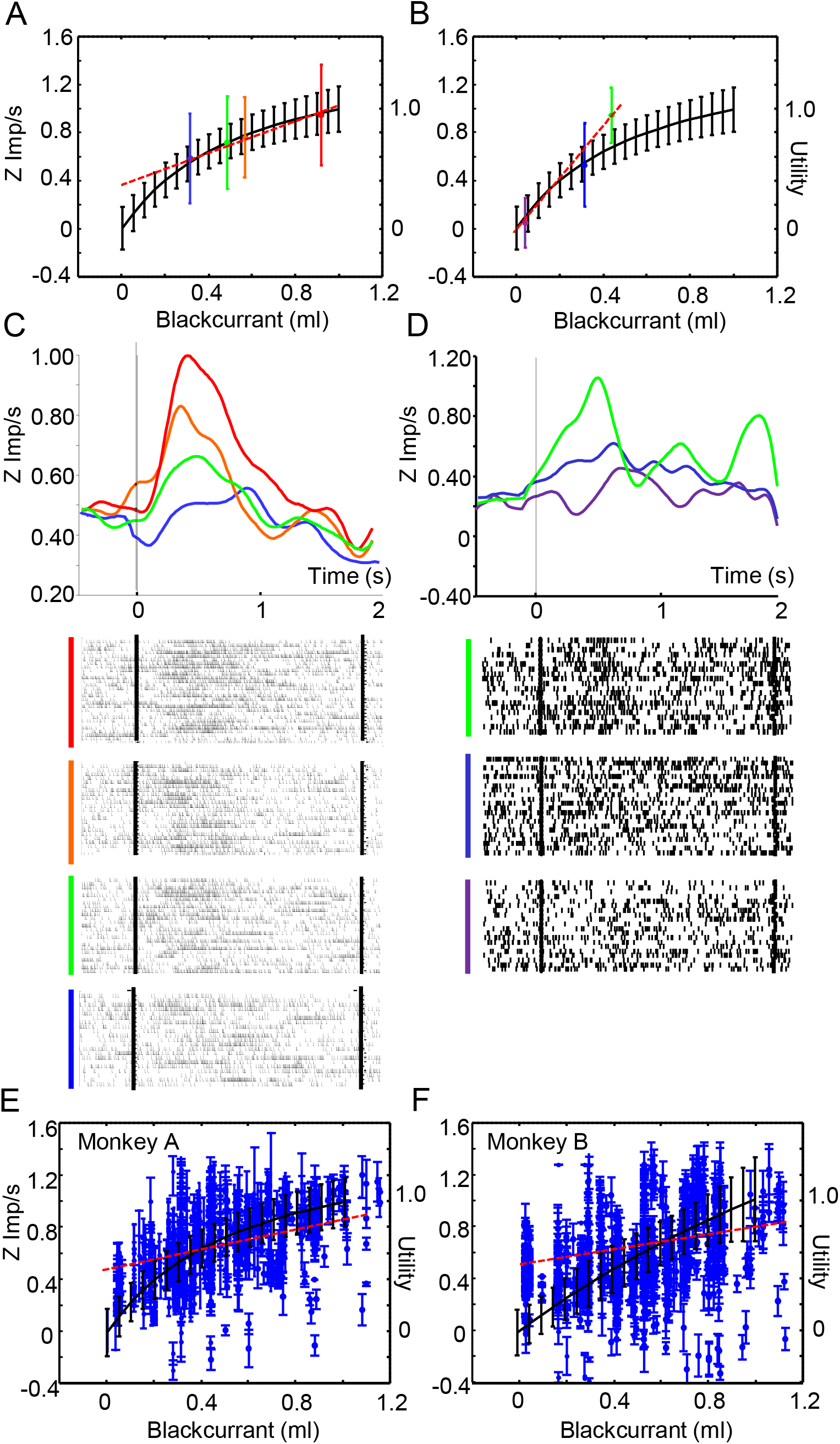
Neuronal coding of utility. (A) and (B) show Z-scored responses (left axis) at target onset for different bundles in two single neurons. The responses and SEM are shown in different colors (red to blue color code, ranging from higher to lower responses). The random utility of the bundles is shown on the right axis. The panels (C), and (D) show response histograms for the two neuron examples along with corresponding rasters. Panels (E) and (F) show the population responses for all neurones at target onset against the random utility of blackcurrant with interaction (black lines) for the bundles blackcurrant vs grape in Subject 1 and blackcurrant vs mango in Subject 2. The population fits with physical value are shown in red (solid line).

Although regressing the example neuron responses with utility (Figure 4C) gave a robust fit (R^2^: 0.9877, p: 0.0062), regressing the same responses with physical value yielded a strongly linear (36 degs) and equally robust fit (R^2^: 0.9956, p: 0.0022). We observed the same effect for the second example neuron in B and D both at Target onset and in other task epochs like Choice and Reward (Figure S3). The regression of neuron response on utility was robust (R^2^: 0.9961, p: 0.0398) but inconclusively similar to the regression with physical value (R^2^ of 0.9979, p: 0.0288) with a linear slope of 69 degs. These results suggest that individual neurons, with different slope fits, might encode separate portions of the utility function linearly, rather than parsing the entire utility range non-linearly.

Figure 4 panels E and F show the Z-scored population responses against physical value and random utility of blackcurrant for separate bundles across all epochs: (E) blackcurrant vs grape (20 neurons, 1800 responses) in Subject 1; and (F) blackcurrant vs mango (31 neurons, 1904 responses) in Subject 2. In these two separate bundles the population the fits with physical value (red lines) were significant but weak (R^2^: 0.035784, p: 1.33 e-4) in comparison with random utility (R^2^: 0.15, p: 6.3e-16). The population responses for each separate bundle and epoch are shown in Figure S4. In the bundle blackcurrant vs grape (A-D), 18 neurons out of 20 (90%, 450 responses) were statistically significant at Target onset. 9 neurons (45%, 213 responses) were significant at GO, 12 neurons (60%, 225 responses) were significant at Choice and 16 neurons (80%, 389 responses) at Reward. Likewise, in the bundle blackcurrant vs water, 13 neurons out of 23 (57%, 349 responses) were statistically significant at Target onset, 12 neurons were significant at GO (52%, 359 responses), 8 neurons at Choice (35%, 262 responses), and 13 neurons (57%, 317) at Reward. These results suggest a stronger coding of utility at target onset followed by choice and reward in Subject 1.

A similar trend was observed in Subject 2 for the bundle blackcurrant vs mango (Figure 4 F) and blackcurrant vs water (Figure S4 G-R). The regression of neuron responses with physical value was weak (R^2^: 0.037567, p: 1.66 e-4) and stronger and highly significant against utility (R^2^:0.18, p: 5.20 e-35). However, for these two bundles the shape of the utility function was also quasi-linear. 20 neurons of out of the 31 significant neurons in the bundle blackcurrant vs mango across all epochs (65%, 476 responses) were statistically significant only at target onset. 13 neurons (42%, 333 responses) were significant at GO, 13 neurons (42%, 260 responses) were significant at Choice and 12 neurons (39%, 309 responses) were significant at Reward.

Table S1-3 and Figure S4 E-F shows the utility slope β and p-value distributions for these bundles. We found that 20 out of 22 neurons (91%, 1800 responses) had statistically significant β coefficients in any epoch for the bundle blackcurrant vs grape (Table S1). Likewise, 31 out of 32 neurons (97%, 1904 responses) had statistically significant utility slope β coefficients in the bundle blackcurrant vs mango (Figure S4 G-L, Table S2) and 23 out of 25 neurons (92%, 1287 responses) were significant in the bundle blackcurrant vs water (Figure S4 M-R, Table S3). Altogether, 74 neurons representing 94% (74 out of 79) of the neurons tested in the fundamental bundles contained statistically significant responses against behaviorally calculated utility in any task epoch. The incorporation of neuronal responses (Table S4) yielded significant improvements in all three representative bundles were electrophysiological data was collected suggesting that neuron responses play a modest but significant contribution to utility and choice prediction.

### Neuronal coding of relative chosen utility

As shown previously (Pastor-Bernier et al., 2019) a significant fraction of neurons represents chosen-value irrespective of any other option (34%, absolute chosen-value coding) or chosen-value coding relative to the unchosen option (12%). Relative chosen-value neurons are of interest because, unlike absolute chosen-value neurons, they directly encode the probability of choice. In our previous study, relative chosen-value was defined indirectly as the neurons that encoded chosen-value or unchosen-value. Presently, we investigated probability coding more directly. We regressed the Z-scored activity for all revealed preference neurons (105) in three bundles and two subjects (79 positive and 26 negative coding neurons) against the probability of choice for either the fixed reference bundle or the variable bundle. 91 neurons had enough data to conduct this double regression analysis (Star Methods). This analysis identified three balanced groups of neurons. 26 neurons (16 positive and 10 negative coding neurons out of 91 tested, 29%) represented the probability in the chosen option irrespective to the alternative bundle and correspond to classic absolute chosen-value coding neurons (Padoa-Schioppa and Assad, 2006). Another 25 neurons (16 positive, 9 negative coding; 28%) represented the probability of the unchosen option and correspond to absolute unchosen-value coding. A third and final group of 26 neurons (15 positive and 11 negative coding; 29%) contained single neurons with statistically significant β coefficients in either choice over the fixed reference bundle or the alternative bundle. The neuron responses in the last group depended on the alternative option both when the fixed reference bundle or the variable bundle were chosen. These 26 neurons constitute good candidates for relative chosen-utility coding. Table S5 summarizes the results for these three categories.

Figure 5 shows single neuron examples of positive and negative relative chosen value neurons. Figure A-B shows the direct correspondence between responses at target onset and choice probability in a positive coding neuron. In (A) the neuron shows a reduction in response with decreasing choice probability on a fixed reference bundle (star) with respect to unchosen alternative options of increasing value (color dots from green to red). Figure 5B shows an increase in response of the neuron with increasing value on the chosen alternative option.

**Figure 5.**
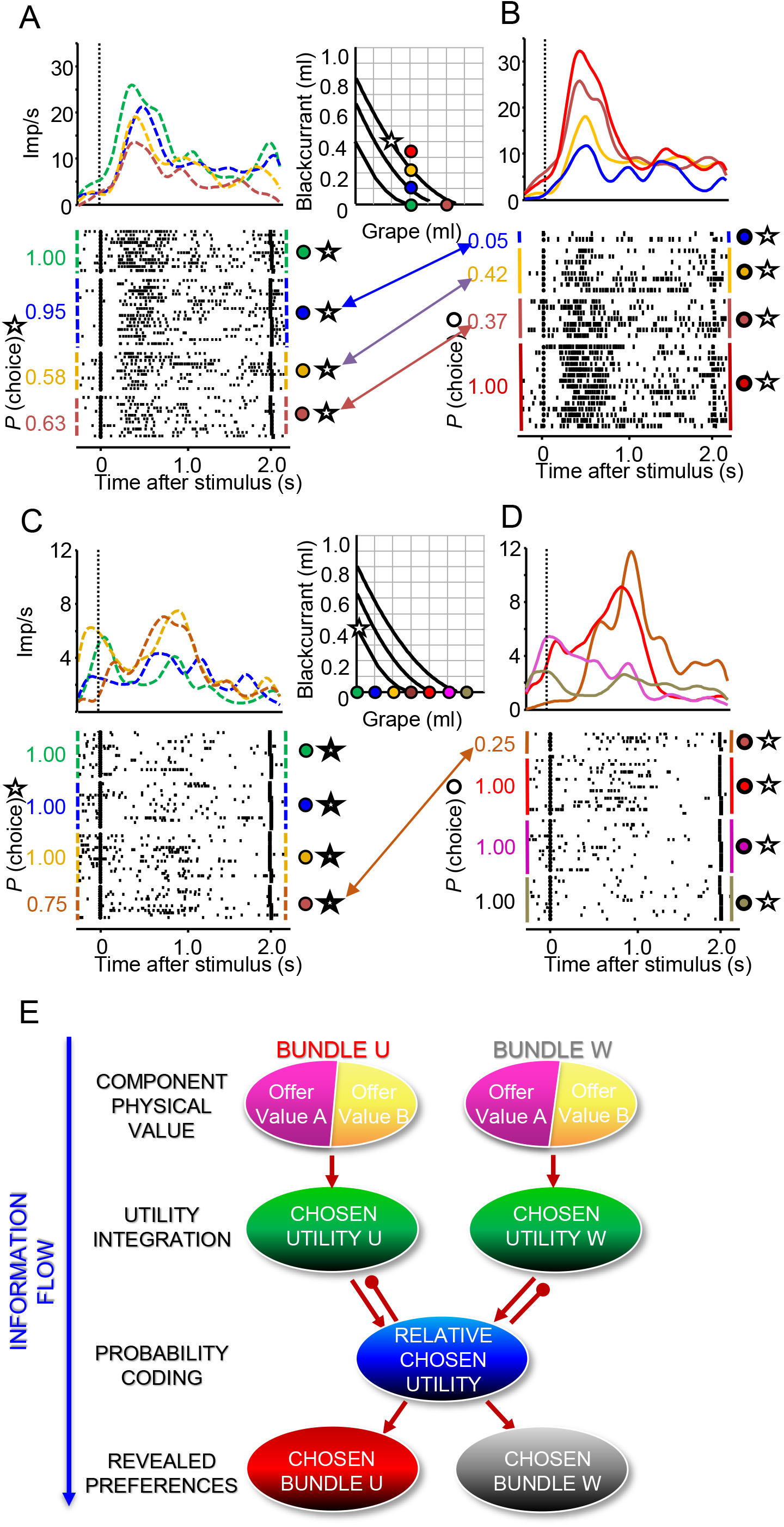
Relative chosen value coding and utility information flow. Single neuron example (A) showing decreasing responses for a chosen constant reference bundle (star) over unchosen alternative bundles of increasing payoff (color dots).The responses for the chosen variable bundles increase monotonically in (B). (C) and (D) shows an example of an inverse coding neurone. The neuron shows monotonically increasing responses for a chosen constant reference bundle against variable bundles of increasing IC and monotonically decreasing responses for chosen variable bundles of increasing value. (E) Information flow suggesting a computational mechanism by which relative chosen value neurones can be instrumental in the specification of revealed preferences in OFC.

Figure 5C-D shows a negative coding neuron with a response at target onset that is inversely proportional to choice probability. The neuron increases response for low choice probability in either the fixed bundle option (C) or variable bundle option (D). Coding of relative chosen value neurons was observed in both subjects (Figure S5) and across several task epochs (Figure S6).

Utility coding depended on the chosen option for these neurons (Figure 6A-F). Positive relative chosen value neurons represented utility only for choices on the variable bundle. Negative relative chosen value neurons represented utility only when the reference bundle was chosen.

**Figure 6.**
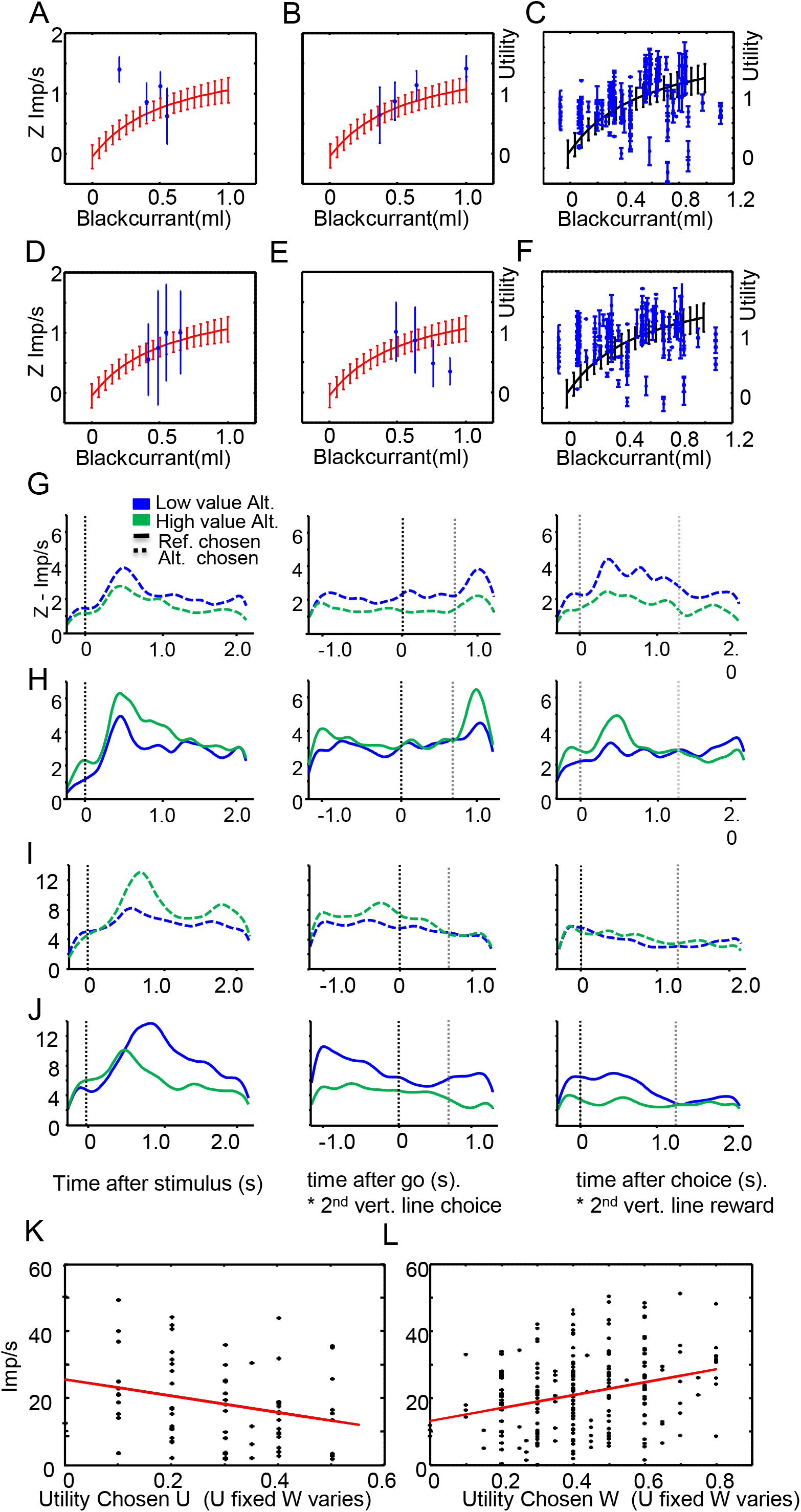
Single neuron and population time-course activity for relative chosen utility. (A) Positive coding neuron showing a decrease of response (blue dots and SEM lines) for the same chosen reference bundle and unchosen alternative bundle of increasing value (B). The same neurone shows an increase in response when the alternative bundles are chosen. The RUM for blackcurrant (red) with random error estimates (vertical lines) is shown. The Z-scored responses for the chosen alternative bundle fall within the range predicted by the RUM for blackcurrant. (D and E) have same convention as above but refer to a negative coding relative chosen-value neuron. (E and F) show RU of z-scored responses for statistically significant utility coding relative-chosen neurones for alternative chosen bundles of increasing value at Target onset and Choice. For the same chosen reference bundle (G), positive-coding relative chosen-value neurones show a reduction in population time-course activity over an alternative high-payoff unchosen option (green), and an increase in response over an alternative low-payoff unchosen option(blue).The same neurones show an increase in response for increasing value in the chosen variable (H) across four task epochs (right panels). Negative coding neurons (I) show an increased population time-course activity over an alternative high-payoff unchosen option and a decreased response for an alternative low-payoff unchosen option. The neuronal responses remains distinctive over Target onset, Go, Choice and Reward epochs(right panels). The same neurones show inverse monotonicity for increasing chosen value in (J). The averaged target-onset single-neurone responses scatter plots for chosen reference bundle (K) or chosen alternative bundle (L) over utility of blackcurrant show a similar pattern of increasing and decreasing monotonicity with choice.

Table S6 summarizes the results for the utility coding transitions along four distinct task epochs, Target onset, GO, Choice and Reward. 15 out of 26 (58%) positive coding neurons represented relative chosen utility and 11 out of 26 (42%) neurons represented chosen utility in negative coding neurons. The positive coding neurons in Table S6 (pink) and negative coding neurons (yellow) were represented more strongly at target onset and choice.

Figure 6 G-J shows time-course responses across the population. The time course of relative chosen-utility was predominant at Target onset and Choice in both positive coding neurons (Figure 6 G-H) and negative coding neurons (Figure 6 I-J). Relative chosen utility in positive coding neurons (Figure 6 K-L) was modest but statistically significant for choices in the variable bundle (R^2^: 0.083031, p: 0.000013) and significant but anticorrelated for choices in the reference bundle (R^2^: 0.05566148, p: 0.003286).

## DISCUSSION

### UTILITY CODING IN OFC

In the current study, we used McFadden’s economic random utility to model behavioral preferences in 10 different bundles and two subjects. The utility functions accounted for the shape of ICs representing the currency and interaction between bundle components. These results suggest that the non-linearity of the indifference maps can derive from the shape of the underlying utility function although this was limited to the set of bundles studied. 54 out of 98 positive coding revealed preference neurons (55%) showed good agreement with behaviourally obtained utility functions suggesting that riskless utility is an important representation for value coding in OFC. Our data had limitations in that we were not able to ascertain whether utility was encoded by the neurons piecewise or whether single neuron coding could span the entire utility range. Our analysis seemed to point that this was the case at population level. Further experiments with a wider titration range of utility and without the constrains that the ordinal characterization of IC imposes would benefit future investigation. In our study an interesting group of 26 neurons, not formally encoding revealed preferences, encoded instead the relative chosen utility of the options. Of those 15 neurons singly encoded the utility of the bundle chosen and 11 neurons encoded only the utility of the alternative unchosen bundle. We suggest a general mechanism incorporating formal utility and a novel relative chosen utility representation for multi-component decision-making. Coding of utility has been previously shown in striatal dopamine neurons (Stauffer et al, 2014) although only from the perspective of either VonNeumann and Morgernstern theory or Kahneman’s economic Prospect theory, and never before in a riskless choice scenario. In the current paper we used McFaddens random utility model and developed the foundations for multi-component and riskless choices in the monkey.

OFC neurons are known to code single rewards, notably offer value coding (Padoa-Schioppa and Assad, 2006). The existence of single component coding neurons was confirmed in our study (N:144: 44% tested neurons in choice over zero bundle) and provided a contrast to revealed preference coding (N:139, 43% of tested neurons in choice over zero bundle). We observed a very balanced proportion of neurons with positive (56%) and negative slope (44%) coding separate components. Competition between different pools of neurons as specified by mutual inhibition between offer-value neurons could be instrumental for choice. Recent evidence (Ballesta and Padoa-Schioppa, 2019) argues against the idea that decisions could be made through mutual inhibition between pools of offer value neurons on the basis that previous beta regressor anticorrelations (Strait et al., 2014; 2015) were confounded by differences in value range. In our study the value range tested was the same for both offers. However, our use of bundle choice options uncovered that a significant fraction of single-reward coding neurons (55%) reflected various forms of chosen value including previously reported absolute chosen value (30%), unchosen value (17%) and relative chosen value (8%) (Pastor-Bernier et al., 2019). In the present study, we investigated neuronal probability coding in 3 different bundles and 2 subjects (105 neurons) and confirmed a balanced presence of absolute chosen utility coding, unchosen utility and relative chosen utility coding in OFC (29%, 28% and 28% respectively). Positive (58%) and negative relative chosen utility neurons were equally present (42%). The information flow scheme depicted in Figure 5E illustrates a transition between valuation and decision processes and suggests a computational mechanism by which relative chosen utility neurons specify revealed preferences in OFC. Although absolute chosen utility is important for value integration and formal economic utility, relative chosen utility neurons directly predict the animal’s choice and are a step closer to the output of the decision process. It is important however to mention that the proposed scheme does not preclude any form of parallel processing, which is known to occur during learning and in other contexts like decisions between actions (Cisek & Pastor-Bernier, 2014). In fact, a temporal analysis of chosen value (Pastor-Bernier et al, 2019) and relative chosen utility in the present work, suggest an early coding of these representations already at Target onset, well before a decision is made. Thus, these representations of economic value, chosen utility and relative chosen utility, might coexist in time rather than succeed each other serially. From a mechanism point of view, inverted negative chosen utility neurons, are also intriguing. Mongillo, Rumpel and Lowenstein, 2018, showed recently that patterns of neural activity in the mammalian brain are primarily determined by inhibitory connectivity, even though most synapses and neurons are excitatory. The percentage of inhibitory neurons required for maximizing performance in many learning and decision-making network models seems to be as high as 30% (Arieli et al., 1996; Capano et al., 2015). This proportion guarantees both the network excitability and the response variability observed. We postulate that the negative coding revealed preference neurons represent inhibitory units that modulate the more widespread positive revealed preference neurons in order to control neuron excitability and guarantee the fidelity of the choice. The proposed scheme also suggests two phases (Figure 5E) involving on one hand utility calculation and on the other hand a comparison of chosen utility alternatives by means reciprocal excitatory and inhibitory projections. According to this scheme, the specification of preferences (choice probability) could be the result of an interaction between neurons encoding relative chosen utility. Our results suggest an important participation of relative chosen utility in revealed preference coding. Future research in this direction would help shed light on the OFC components involved in value-to-choice transformations.

## Acknowledgements

We thank Chrissy Thompson for invaluable help with animal training, Paul Cisek for sharing his SQL-Matlab toolbox (NeuroMath), Charles R. Plott, Christopher Harris, Simone Ferrari-Toniolo and Fabian Grabenhorst for inspiration and insightful comments on experimental economics and neuronal data analysis. The Wellcome Trust, European Research Council (ERC) and US National Institutes of Mental Health (NIMH) Caltech Conte Center supported this work.

## Author contributions

A.P.-B. and W.S. designed the research, A.P.-B. performed the experiments, A.P.-B. analysed the data, A.P.-B. and W.S. wrote the manuscript.

## Competing Interests

The authors declare no competing interests.

---

**SUPPLEMENTARY TABLE 1.** RUM regression coefficients and p-Values for individual neurons in the bundle blackcurrant vs grape and subject 1

| Neuron | Target Onset |  | GO |  | Choice |  | Reward |  |
| --- | --- | --- | --- | --- | --- | --- | --- | --- |
| | Utility $\beta$ | pValue | Utility $\beta$ | pValue | Utility $\beta$ | pValue | Utility $\beta$ | pValue |
| 1 | 0.77 | 1.00E-02 | 0.81 | 6.00E-02 | 0.46 | 3.50E-01 | -1.06 | 0.48 |
| 2 | -0.18 | 6.00E-01 | 1.01 | 1.30E-01 | 1.71 | 1.00E-02 | 2.02 | 5.00E-03 |
| 3 | 0.26 | 3.20E-01 | 0.4 | 2.60E-01 | 0.1 | 4.70E-01 | 0.9 | 5.00E-03 |
| 4 | 1.93 | 1.00E-02 | 1.71 | 1.00E-02 | 1.78 | 5.00E-03 | 1.19 | 1.00E-02 |
| 5 | 0.34 | 5.00E-03 | 0.57 | 5.00E-03 | 0.52 | 2.00E-02 | 0.37 | 2.00E-02 |
| 6 | 1.65 | 5.00E-03 | 1.22 | 1.00E-02 | 1.42 | 1.00E-02 | 0.7 | 5.00E-03 |
| 7 | 1.07 | 2.30E-01 | 2.22 | 3.00E-02 | 1.67 | 5.00E-02 | 0.88 | 2.00E-01 |
| 8 | 1.27 | 3.00E-02 | 2.3 | 1.00E-02 | 1.57 | 4.00E-02 | 0.98 | 1.30E-01 |
| 9 | 1.54 | 4.00E-02 | 1.15 | 1.00E-01 | 0.79 | 2.30E-01 | 1.52 | 2.00E-02 |
| 10 | 0.58 | 3.00E-02 | 0.11 | 6.10E-01 | -0.1 | 6.00E-01 | -0.16 | 5.50E-01 |
| 11 | 0.15 | 2.00E-02 | 0.17 | 4.00E-02 | 0.2 | 1.00E-02 | 0.09 | 2.80E-01 |
| 12 | 1.63 | 1.00E-02 | 2.49 | 5.00E-03 | 3.27 | 5.00E-03 | 1.69 | 2.00E-02 |
| 13 | 0.09 | 5.00E-03 | 0.04 | 3.80E-01 | 0.07 | 9.00E-02 | 0.02 | 6.00E-01 |
| 14 | 0.06 | 3.20E-01 | 0.41 | 5.00E-02 | 0.4 | 5.00E-03 | 0.29 | 5.00E-03 |
| 15 | 0.51 | 1.20E-01 | 0.92 | 2.00E-02 | 0.93 | 2.00E-02 | 0.4 | 2.20E-01 |
| 16 | 0.14 | 5.00E-03 | 0.21 | 5.00E-03 | 0.22 | 5.00E-03 | 0.12 | 5.00E-03 |
| 17 | 2.51 | 5.00E-03 | 2.79 | 5.00E-03 | 3.3 | 5.00E-03 | 1.57 | 4.00E-02 |
| 18 | 0.57 | 6.00E-02 | 0.72 | 5.00E-03 | 0.93 | 5.00E-03 | 0.83 | 1.00E-02 |
| 19 | 2.25 | 5.00E-03 | 2.13 | 5.00E-03 | 2.39 | 5.00E-03 | 1.01 | 6.00E-02 |
| 20 | 0.29 | 2.50E-01 | 0.23 | 3.30E-01 | 0.39 | 1.00E-01 | 0.68 | 5.00E-03 |

**SUPPLEMENTARY TABLE 2.** RUM regression coefficients and p-Values for individual neurons in the bundle blackcurrant vs mango and subject 2

| Neuron | Target Onset |  | GO |  | Choice |  | Reward |  |
| --- | --- | --- | --- | --- | --- | --- | --- | --- |
| | Utility $\beta$ | pValue | Utility $\beta$ | pValue | Utility $\beta$ | pValue | Utility $\beta$ | pValue |
| 1 | 0.58 | 3.00E-02 | 0.35 | 2.00E-01 | 0.39 | 1.80E-01 | 0.23 | 6.10E-01 |
| 2 | 0.61 | 1.40E-01 | 0.8 | 5.00E-02 | 0.74 | 9.00E-02 | 1.56 | 5.00E-03 |
| 3 | 0.01 | 4.00E-02 | 0.01 | 4.00E-02 | 0.01 | 4.00E-02 | 0.01 | 4.00E-02 |
| 4 | -0.05 | 1.30E-01 | 0.01 | 9.80E-01 | 0.04 | 4.00E-02 | 0.08 | 4.40E-01 |
| 5 | 0.42 | 2.00E-02 | 0.02 | 9.10E-01 | 0.04 | 8.00E-01 | 0.02 | 9.10E-01 |
| 6 | 0.61 | 5.00E-03 | 0.09 | 7.00E-01 | -0.09 | 6.70E-01 | -0.17 | 3.50E-01 |
| 7 | 0.08 | 2.00E-02 | 0.04 | 5.00E-01 | 0.02 | 5.40E-01 | 0.04 | 1.70E-01 |
| 8 | 0.14 | 6.00E-02 | 0.21 | 7.00E-02 | 0.26 | 5.00E-02 | 0.23 | 1.70E-01 |
| 9 | 0.61 | 1.40E-01 | 0.8 | 5.00E-02 | 0.74 | 9.00E-02 | 1.56 | 5.00E-03 |
| 10 | 1.54 | 4.00E-02 | 1.9 | 8.00E-02 | 3.41 | 2.00E-02 | 1.41 | 2.00E-02 |
| 11 | 0.46 | 2.70E-01 | 1.51 | 5.00E-02 | 1.11 | 1.20E-01 | 0.84 | 2.80E-01 |
| 12 | 0.97 | 5.00E-03 | -0.4 | 5.00E-03 | -0.65 | 1.00E-01 | -0.98 | 1.80E-01 |
| 13 | 4.55 | 2.00E-02 | -0.7 | 2.10E-01 | -0.52 | 2.20E-01 | -0.53 | 3.10E-01 |
| 14 | 1.24 | 4.00E-02 | 1.32 | 8.00E-02 | 0.88 | 1.40E-01 | 0.23 | 6.50E-01 |
| 15 | 0.72 | 6.00E-02 | 1.37 | 1.00E-02 | 1.44 | 5.00E-03 | 0.49 | 3.40E-01 |
| 16 | 0.4 | 2.00E-02 | 0.5 | 1.80E-01 | 0.04 | 8.80E-01 | 0.01 | 9.90E-01 |
| 17 | 0.59 | 3.00E-02 | 0.64 | 3.00E-02 | 0.58 | 1.00E-02 | 1.07 | 3.00E-02 |
| 18 | 0.6 | 5.00E-02 | 0.38 | 2.60E-01 | 0.91 | 2.00E-02 | 0.29 | 3.90E-01 |
| 19 | 1.3 | 6.00E-02 | 0.07 | 9.00E-01 | 1.2 | 2.30E-01 | 1.71 | 2.00E-02 |
| 20 | 1.12 | 4.00E-02 | 0.6 | 4.00E-01 | 0.13 | 8.90E-01 | 1.78 | 4.00E-02 |
| 21 | 0.6 | 1.00E-02 | 0.6 | 3.00E-02 | 1.02 | 5.00E-03 | 0.69 | 1.50E-01 |
| 22 | -0.06 | 3.00E-01 | -0.07 | 2.10E-01 | 0.03 | 3.80E-01 | 0.15 | 5.00E-03 |
| 23 | -0.16 | 4.50E-01 | -0.21 | 4.50E-01 | 0.36 | 9.00E-02 | 1.28 | 5.00E-03 |
| 24 | 3.05 | 5.00E-03 | 3.42 | 3.00E-02 | 2.39 | 6.00E-02 | 2.59 | 5.00E-02 |
| 25 | 0.35 | 4.00E-02 | 0.21 | 7.20E-01 | 0.27 | 7.50E-01 | 0.13 | 8.20E-01 |
| 26 | 3.17 | 1.10E-01 | 2.14 | 1.20E-01 | 2.76 | 2.00E-02 | 0.96 | 4.00E-01 |
| 27 | 1.62 | 5.00E-03 | 0.78 | 5.00E-03 | 1.08 | 5.00E-03 | 1.2 | 5.00E-03 |
| 28 | 1.77 | 5.00E-03 | 1.05 | 5.00E-03 | 1.23 | 5.00E-03 | 1.34 | 5.00E-03 |
| 29 | 0.28 | 3.50E-01 | 0.46 | 2.20E-01 | 0.38 | 3.00E-01 | 1.21 | 3.00E-02 |
| 30 | -0.01 | 8.10E-01 | 0.2 | 2.00E-02 | 0.25 | 5.00E-03 | 0.29 | 1.00E-02 |
| 31 | 0.04 | 5.80E-01 | -0.09 | 2.20E-01 | -0.01 | 9.40E-01 | 0.16 | 3.00E-02 |

**SUPPLEMENTARY TABLE 3.**
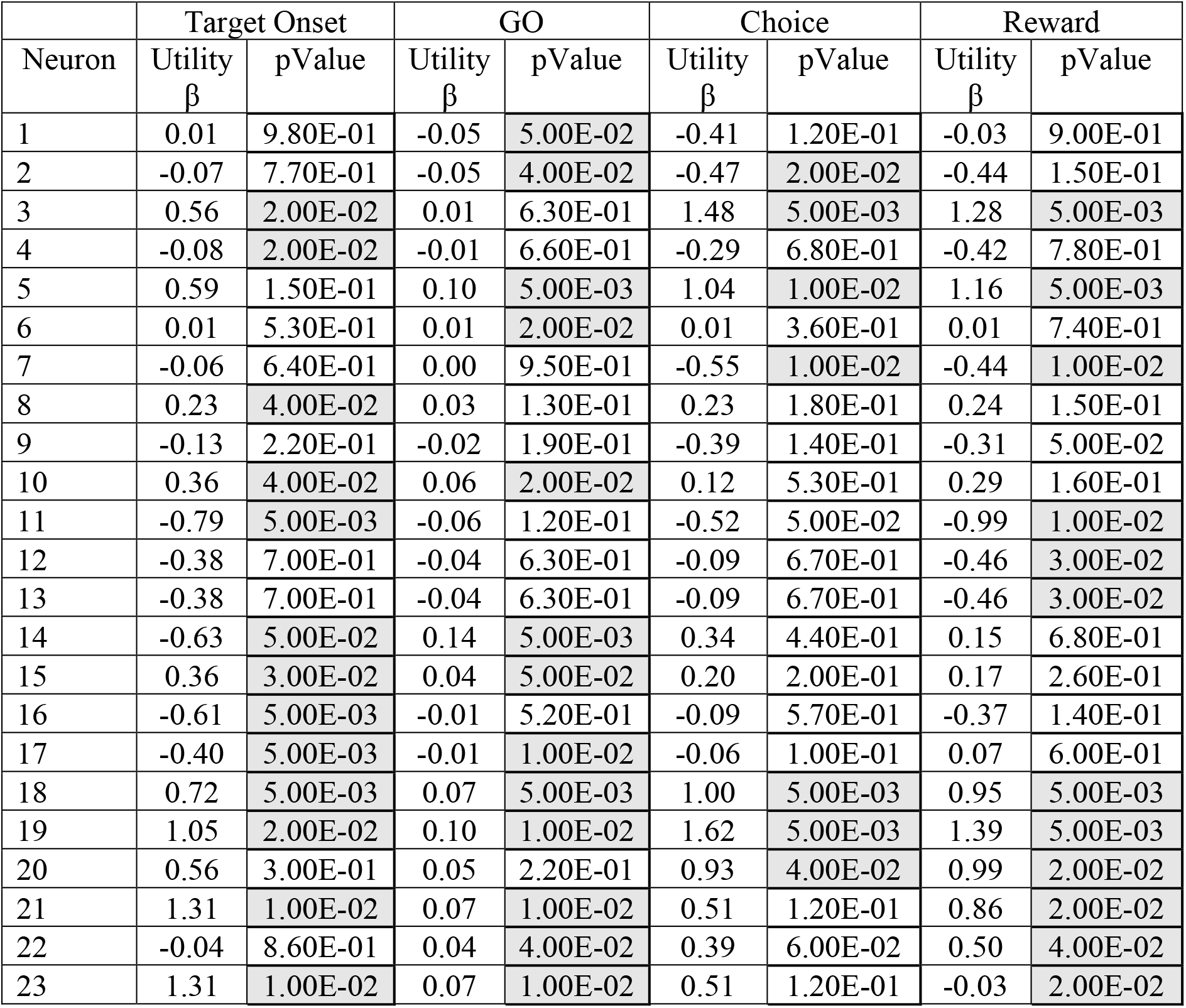
RUM regression coefficients and p-Values for individual neurons in the bundle blackcurrant vs water and subject 2

**SUPPLEMENTARY TABLE 4.** Neuronal vs Behavioural RUM comparisons

| Bundle Component | Neuronal RUM |  |  | Behavioral RUM |  |  |
| --- | --- | --- | --- | --- | --- | --- |
|  | Log-Likelihood | McFadden R<sup>2</sup> | AIC | Log-Likelihood | McFadden R<sup>2</sup> | AIC |
| Blackcurrant | -662.97 | 0.73072 | 1341.942 | -683.75 | 0.72228 | 1379.504 |
| Grape | -180.94 | 0.66033 | 377.888 | -187.42 | 0.64824 | 386.8373 |
| Water | -1205.8 | 0.55029 | 2427.55 | -1217.4 | 0.54595 | 2446.829 |
| Blackcurrant S2 | -479.92 | 0.60502 | 975.845 | -480.53 | 0.60512 | 973.0554 |
| Mango S2 | -662.97 | 0.73072 | 1341.942 | -683.75 | 0.72228 | 1379.504 |

**SUPPLEMENTARY TABLE 5.**
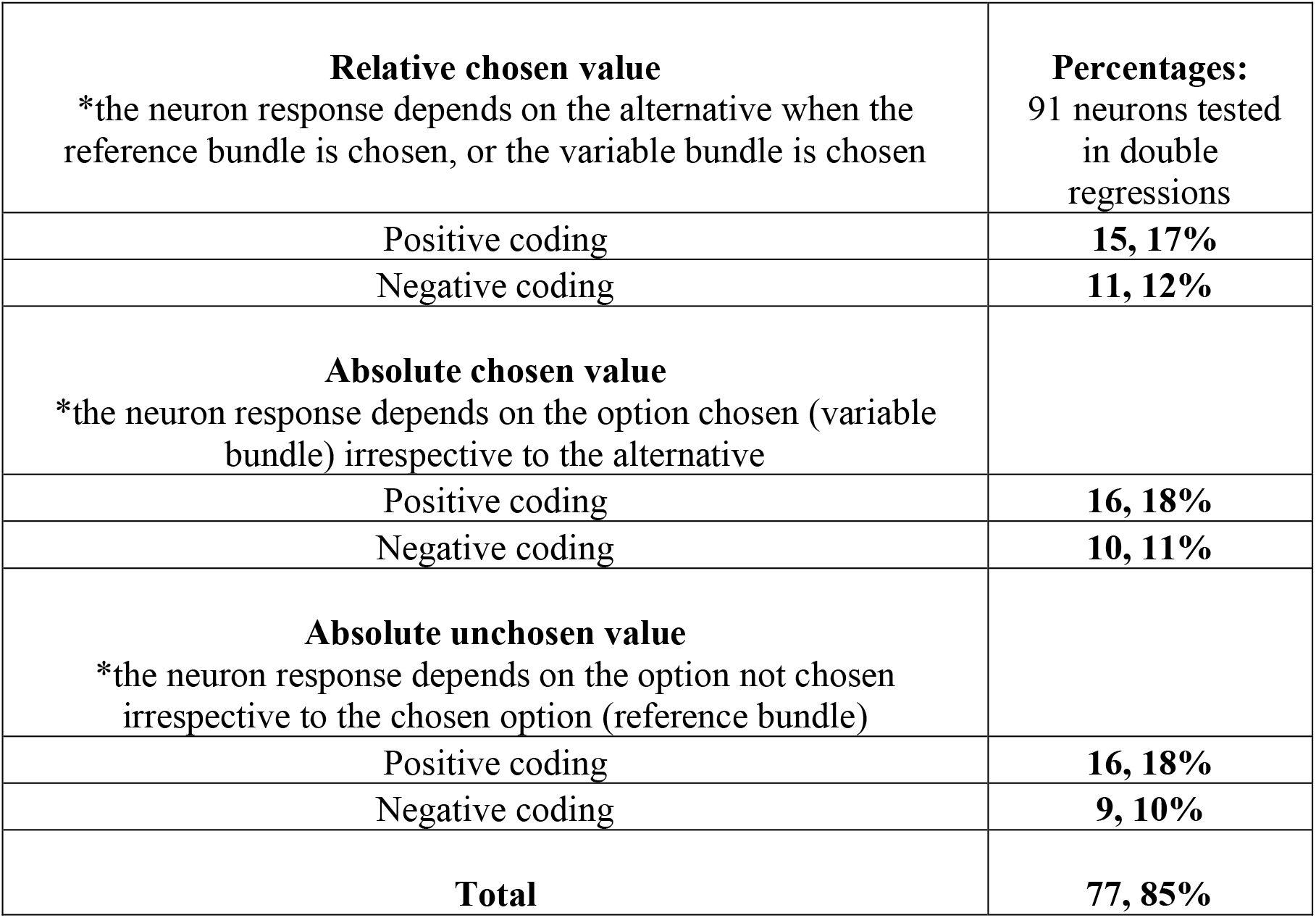
Neuron categories as determined by sign and significance of probability slope coefficients

|  |  |
| --- | --- |
| **Relative chosen value**<br>*the neuron response depends on the alternative when the reference bundle is chosen, or the variable bundle is chosen | **Percentages:**<br>91 neurons tested in double regressions |
| Positive coding | **15, 17%** |
| Negative coding | **11, 12%** |
| **Absolute chosen value**<br>*the neuron response depends on the option chosen (variable bundle) irrespective to the alternative |  |
| Positive coding | **16, 18%** |
| Negative coding | **10, 11%** |
| **Absolute unchosen value**<br>*the neuron response depends on the option not chosen irrespective to the chosen option (reference bundle) |  |
| Positive coding | **16, 18%** |
| Negative coding | **9, 10%** |
| **Total** | **77, 85%** |

**SUPPLEMENTARY TABLE 6.**
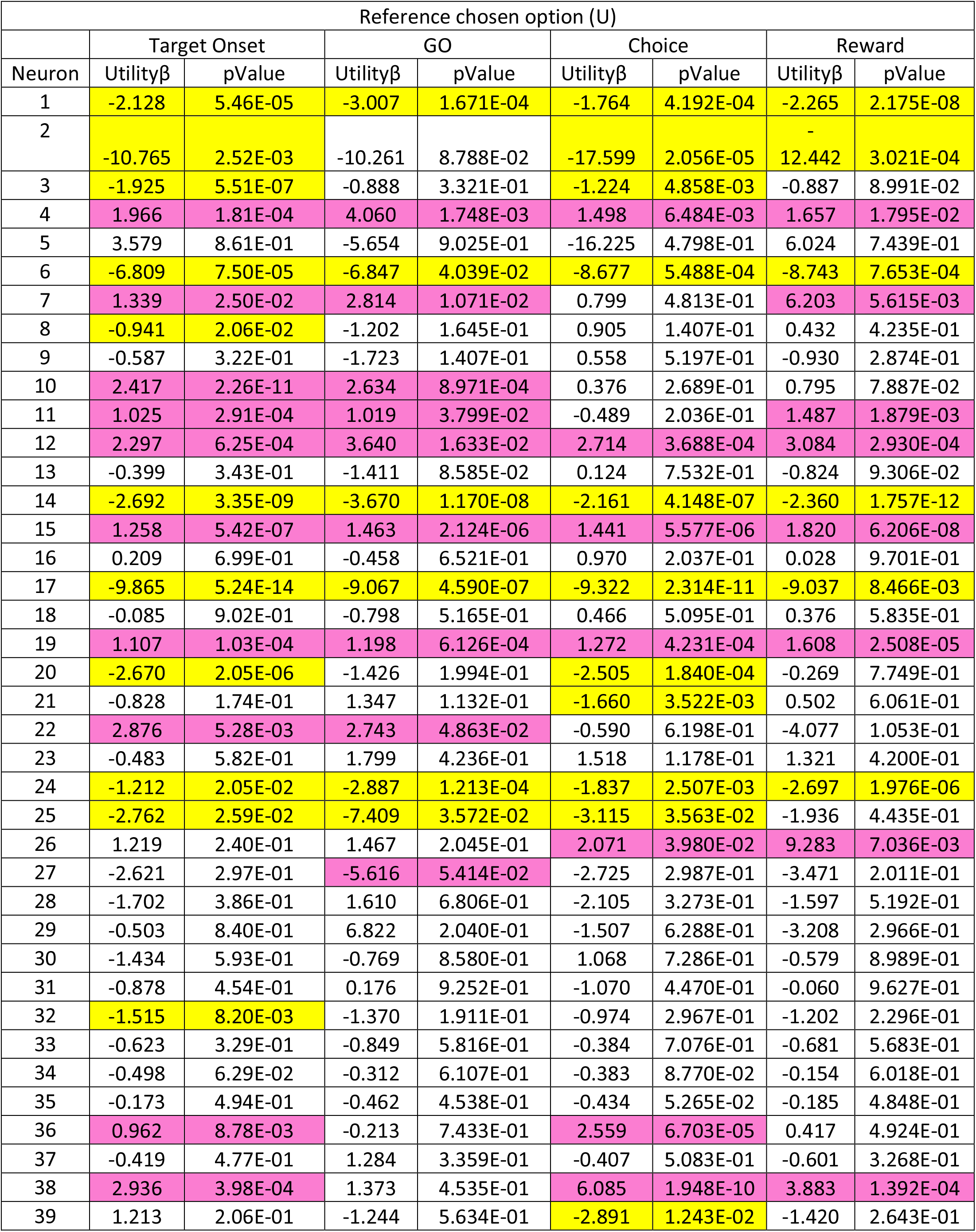
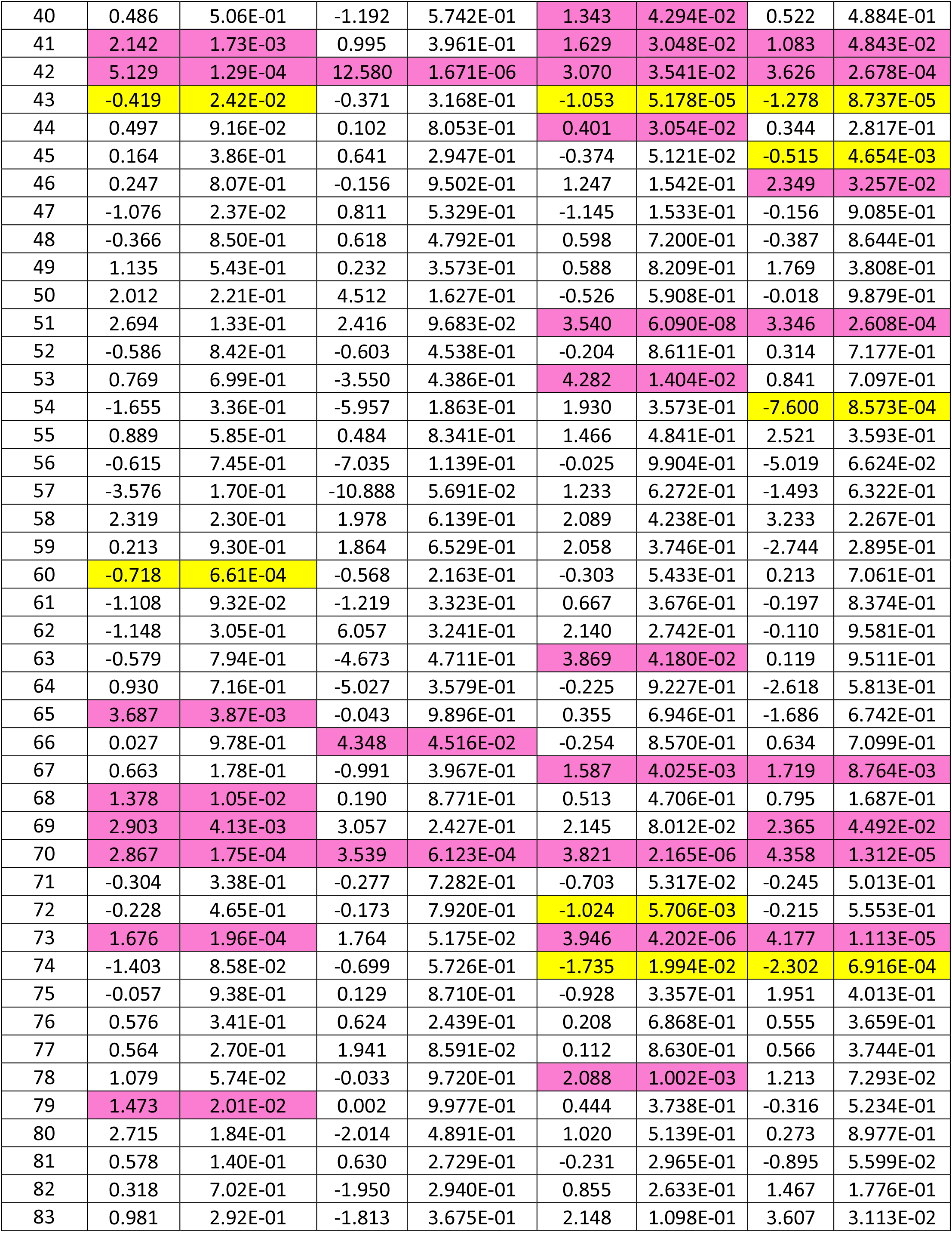
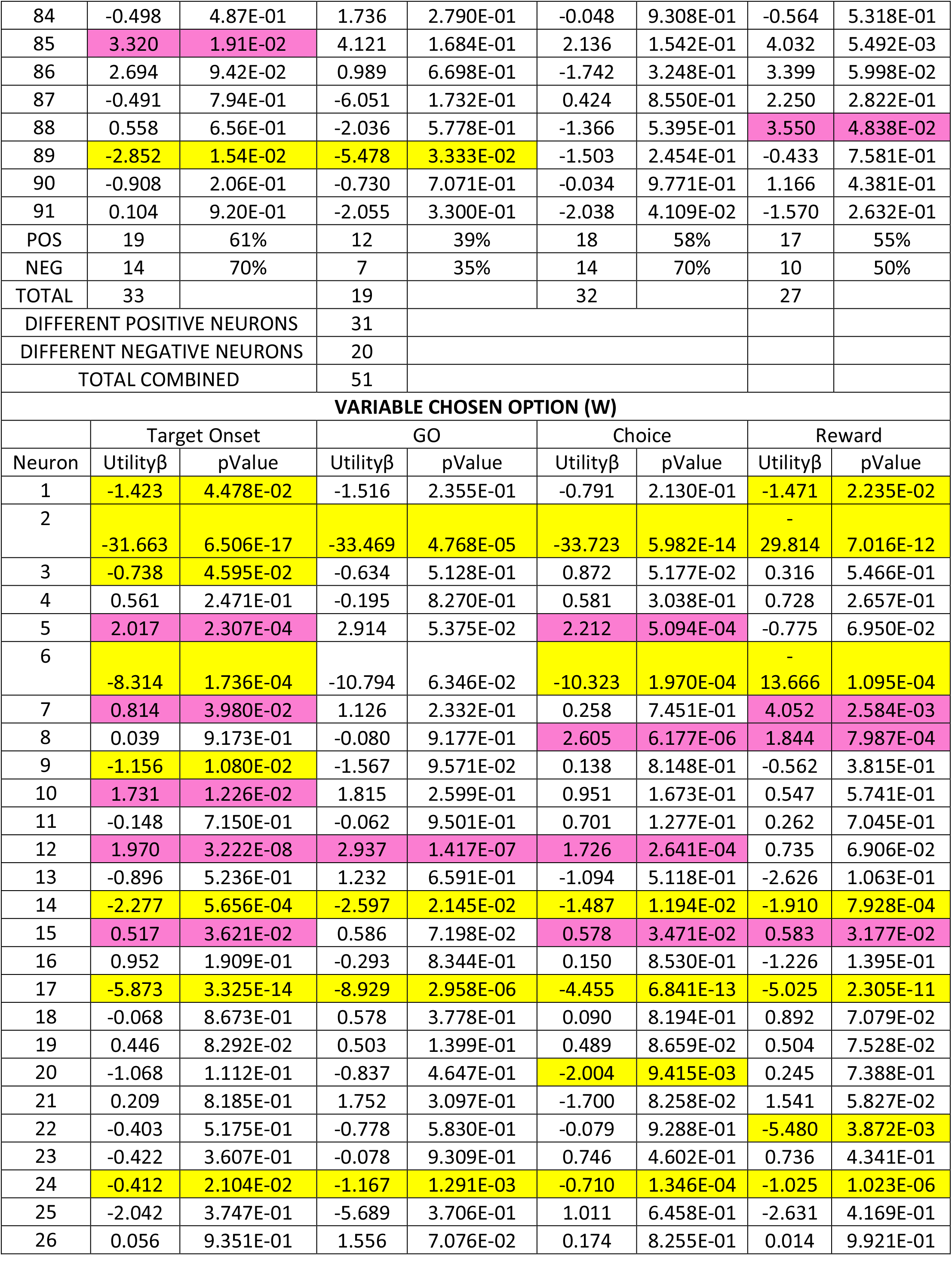
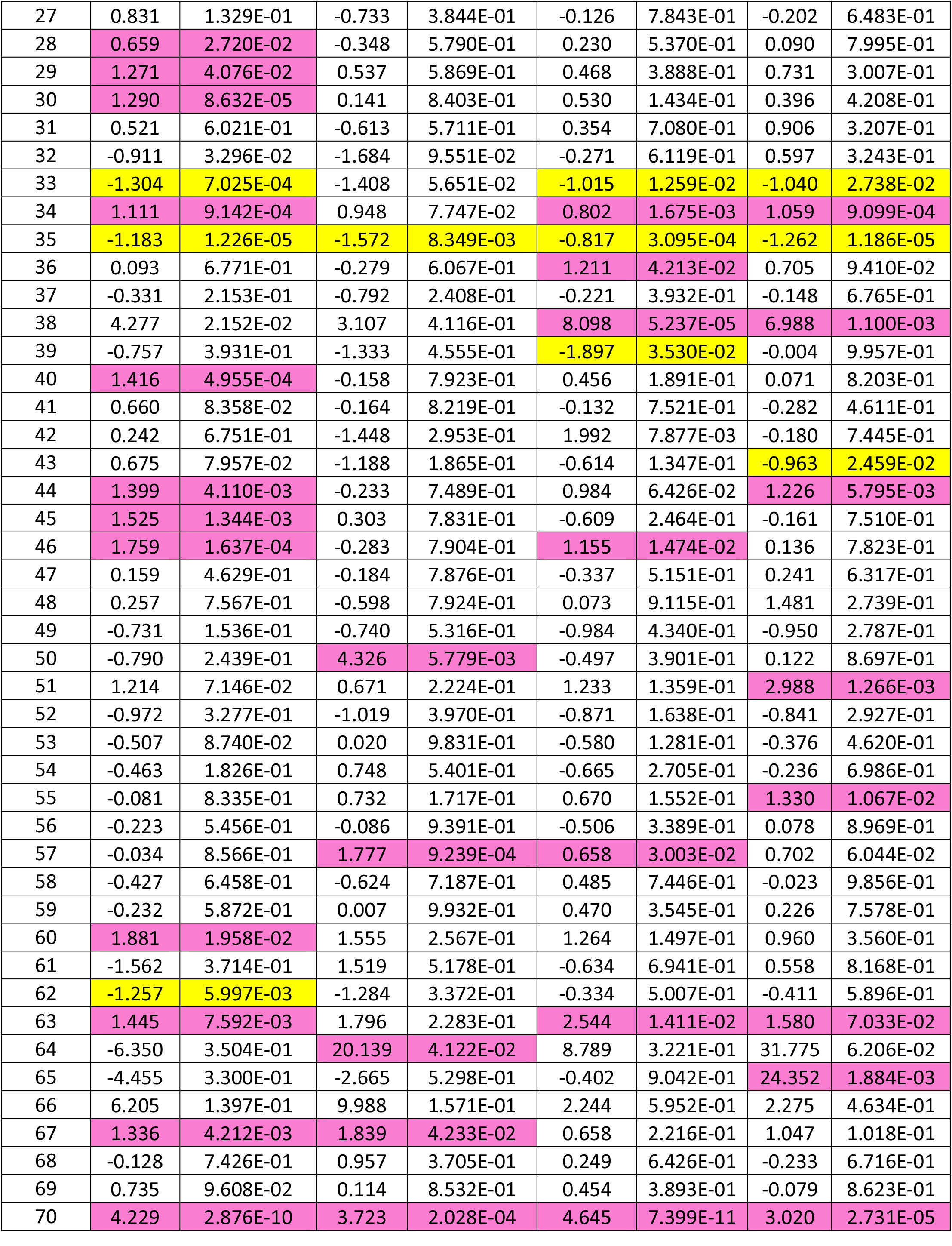
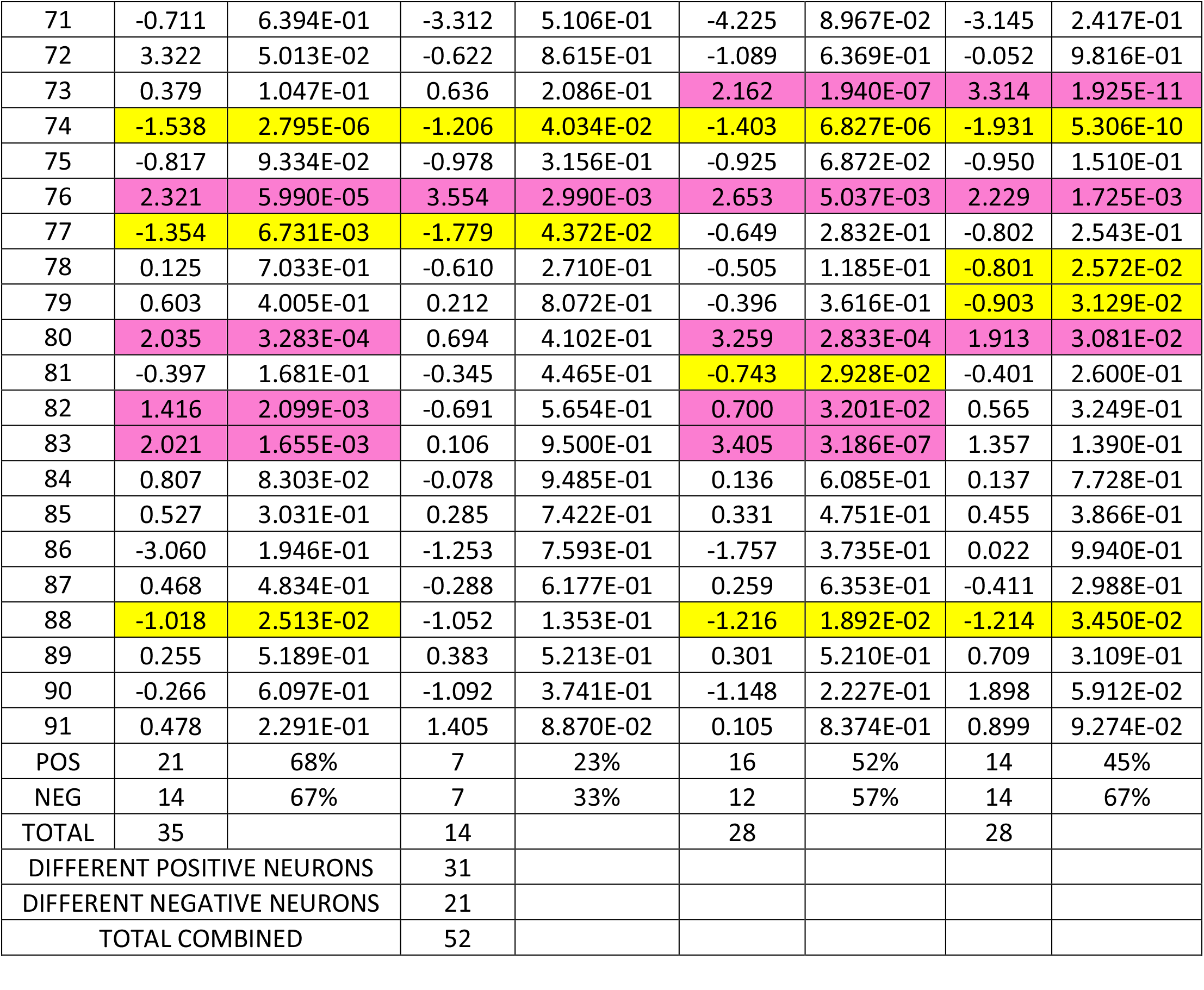
Relativity utility coding

## STAR Methods for

### (S0) Ethics and animals

This research has been regulated, ethically reviewed and supervised by the following UK and University of Cambridge (UCam) institutions and individuals:

- UK Home Office, implementing the Animals (Scientific Procedures) Act 1986, Amendment Regulations 2012, and represented by the local UK Home Office Inspector
- UK Animals in Science Committee
- UCam Animal Welfare and Ethical Review Body (AWERB)
- UK National Centre for Replacement, Refinement and Reduction of Animal Experiments (NC3Rs)
- UCam Biomedical Service (UBS) Certificate Holder
- UCam Welfare Officer
- UCam Governance and Strategy Committee
- UCam Named Veterinary Surgeon (NVS)
- UCam Named Animal Care and Welfare Officer (NACWO)

Two adult male macaque monkeys (*Macaca mulatta*; monkey A, monkey B), weighing 11.0 kg and 10.0 kg, respectively, were used in the experiments. Neither animal had been used in any prior study.

### (S1) General behavior

The animals were habituated during several months to work on a primate chair (Crist Instruments) for a few hours each working day. They were trained in a specific, computer-controlled behavioral task in which they contacted visual stimuli on a horizontally mounted touch-sensitive computer monitor (Elo) located 30cm in front of them. The subjects eye positions in the horizontal and vertical plane were monitored with a non-invasive infrared oculometer (Iscan). Matlab software (Mathworks) running on a Microsoft Windows XP computer controlled the behavior and collected, analyzed and presented data on-line. A solenoid valve (ASCO, SCB262C068) controlled by the same Windows computer served to deliver specific quantities of liquids. A Microsoft SQL Server 2008 Database served for Matlab off-line data analysis. Following task training for 6 months, animals were surgically implanted with a recording chamber for electrophysiological recordings, which typically lasted for another 6-10 months.

### (S2) Visual stimuli and two-component bundles

The computer touch monitor presented the animal with two visual stimuli at the left and right (angle of 4°) representing two bundles, respectively (Figs. 1). Each stimulus indicated a bundle that contained the same two distinct liquid rewards (equivalent to ‘goods’ in economics) with independently set quantities (component A, plotted along the y-axis of a 2D graph, and reward B, plotted along the x-axis). Each reward was presented by a distinctly colored stimulus with a superimposed rectangle containing a value bar whose vertical position indicated the quantity of that. In the standard option set, two bundles were presented at pseudorandomly alternating left and right positions on a computer monitor in front of the animal; each bundle contained two stimuli. In both bundles, component A (top, violet) was always blackcurrant juice, whereas component B (bottom, green) could be grape, water, apple, mango, strawberry, peach, lemon and saline. A fifth bundle type contained monosodium glutamate (MSG) added to blackcurrant juice (component A) and inositol monophosphate (IMP) added to grape juice (component B); this bundle composition was used to test synergistic, enhancing effects of the combination of MSG with IMP.

### (S3) Behavioral task

After the animal’s hand contacted a capacitive touch key, two visual bundle stimuli appeared on the touch monitor (Fig. S1). The left and right positions of the bundle stimuli alternated pseudorandomly, but there was no further distinction between the stimuli that could serve for identifying them as distinct objects. The animal held the touch key for 0-2 s (0-3 s for three bundles), and then a Go signal appeared (blue dots underneath the bundles). Without any further imposed delay, the animal released the touch key after an average reaction time of 445 ms (Monkey A: 396 ms, Monkey B: 493 ms) and touched the blue Go spot underneath the bundle of its choice. This choice revealed the animal’s preference at this moment. The animal kept touching the Go spot for 1 s, after which it received the reward quantities of the chosen bundle, consisting first of reward A, always followed 500 ms later by reward B. By always delivering the reward components in the same sequence, and by always using blackcurrant juice as reward A, we incurred a consistent discount of reward B that contributed a constant, non-varying, constituent factor for the revealed preference for that reward. This delay, rather than a simultaneous delivery of both rewards, was introduced to reduce the interaction between the different reward components of the studied bundles. With two bundle options, the animal chose between a reference bundle (U) and a variable bundle (W) in multiple trials.

We set one reward in the variable bundle to a constant value while pseudo-randomly varying the quantity of the other reward we are wishing to titrate. The variation of the animal’s repeated choice with that single varying reward allowed us to construct a full psychophysical function and estimate the choice indifference point (IP) with Weibull fit (*P* = 0.5 choice of each bundle). We obtained each IP from a total of 80 trials (2 left-right stimulus positions with 5 equally spaced reward quantities in 8 trials). To avoid known adaptations in OFC neurons (Padoa-Schioppa, 2009) we always tested the full reward range of the experiment.

### (S4) Behavioral assessment of indifference curves

The behavioral method used to obtain indifference curves (IC) from choice indifference points has been presented in full detail (Pastor-Bernier et al., 2017). Figure 1 (C-D) illustrates the titration procedure in detail. To obtain an IC, we fit a series of IPs with a hyperbolic function using weighted least mean squares.

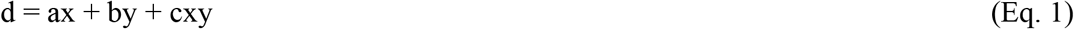

with x and y as reward quantities, a as slope (currency), c as curvature. The hyperbolic function provided the slope (coefficient b) and curvature (coefficient a) of the behavioral IC.

The hyperbolic function can be written in an equivalent form to the regression with interaction used for analysing neuronal responses (Pastor-Bernier et al., 2019)

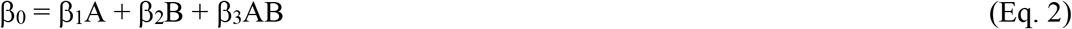

### (S5) Definition and requirement criteria for economic-stable conditions

Satiety was detected by psychophysical choice functions exceeding the confidence intervals of initial tests; this measure indicated changed currency relations between the two bundle rewards. More specifically, the gradual effect of satiety on choice preference was identified by tracking the indifference points (IPs) as consumption advanced across blocks of 80 trials each. The IPs were obtained psychophysically (Weibull fit) for fixed and equally spaced amounts of Juice B. Currency changes were assessed with interleaved anchor trials distributed on identically repeated currency control blocks consisting of choice bundles between the intercept Y axis component blackcurrant and the X axis variable component grape, grape MSG/IMP, water, strawberry, mango, apple, peach, lemon and saline. The IPs were classified as belonging to the sated condition if they exceeded the confidence interval (green shade) defined by the first titrated IP on a currency block as shown previously (Pastor-Bernier et al., 2019). Only data collected in pre-sated and stable economic conditions is included in this paper.

### (S6) Random utility estimation

Random utility models have become a reference approach in economics when one wants to analyse the choice by a decision maker of one among a set of mutually exclusive alternatives. Since the seminal works of Mc Fadden (McFadden 1974) a large amount of theoretical and empirical literature have been developed (Train and Kenneth, 2009).Utility models rely on the hypothesis that the decision maker is able to rank the different alternatives by an order of preference represented by a utility function, the chosen alternative being the one which is associated with the highest level of utility. In random utility models (RUM) part of the utility consists of two parts: a deterministic portion that depends on the differences in the attributes (marginal utilities), and a random deviate which contains all the unobserved determinants of the utility (McFadden, 1973). The random element comes in our case from varying preferences across sessions within a given bundle and subject, which are due to uncontrollable factors such as changes in motivation and internal state.

### (S7) RUM notation

In a general notation, the Utility for each option j within a set of different alternatives i, is given by:

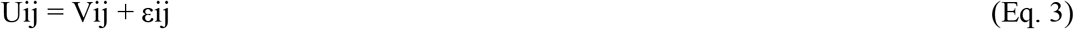

Vij represents the deterministic portion of the utility, and εij the stochastic portion of the utility.

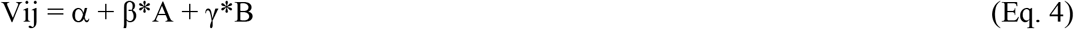

where α is an alternative specific constant, β is the marginal utility of component A (Blackcurrant), and γ is the marginal utility of component B (grape, grape-IMP, strawberry, water, mango, apple, yoghurt or saline). Marginal utilities are estimated using choice data.

The probability of choosing option j among n alternatives is given by

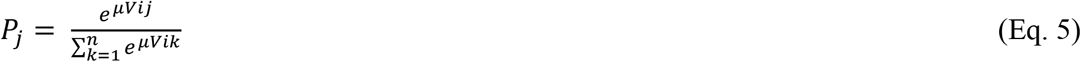

μ is a softmax scale term which is inversely related to the error of the variance, and is typically = 1, unless we wish to incorporate temporal covariates of utility (Cerreia-Vioglio et al., 2017). The relative choice probabilities between options is simply explained by the difference in utility between the options.

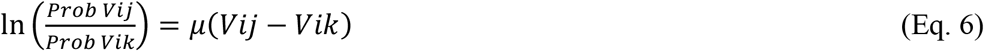

In this framework, any variable that is constant across choices drops out of the probability formula (Eg. Monkey id). Any variable which is constant across alternatives therefore cannot explain the relative likelihood of choice. The only way to let set-specific variables affect choice is to allow the utility function to depend on an interaction variable of interest, like days, sessions or blocks. Combining data for a given bundle and subject across sessions produces a data set with slight preference variations which enables readily estimation of εij, the random error associated with the stochastic portion of the utility function.

### (S8) RUM error generative process

The stochastic portion of the utility εij, captures the preference variability across sessions. Given explicit values of utility for each option j, Vij, where (j=1…n), independent samples are taken randomly for each j across a set i. Thus, Random utility generates a distribution of linearly ordered preferences, such that the conditional probability of a permutation ‘π’ given explicit values of utility Vij produces a distribution in ordered preferences from j=1…n.

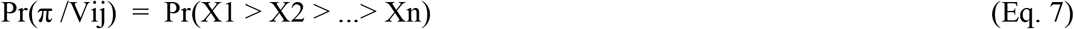

The generative process has been described elsewhere in detail (Parkes et al., 2018). Different hypothesis on the distribution of this random deviate lead to different flavours of random utility models. Early developments of these models were based on the hypothesis of identically and independent errors following a Gumbel distribution, leading to the multinomial logit model (MNL). Individual heterogeneity can be introduced in the parameters associated with the covariates entering the observable part of the utility. This leads respectively to the multinomial mixed effect model (MXL) which is applied here. The multinomial ‘mlogit’ package in R statistical software allows the estimation of these random utility models quite concisely (See Sarrias and Daziano 2017, for a survey of relevant computational packages in R language). mlogit provides a wide set of estimators for random utility with compatible syntax and excellent translation to other analytical software such as Matlab (Mathworks) and Python, which are extensively used in different stages of the present work.

### (S9) McFadden’s maximum likelihood estimation

Logistic regression models are fitted using the method of maximum likelihood - i.e. the parameter estimates are those values which maximize the likelihood of the data which have been observed. McFadden’s R squared measure is defined as:

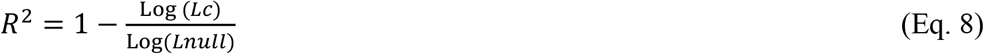

where *Lc* denotes the maximized likelihood value from the current fitted model, and *Lnull* denotes the corresponding value but for the null model - the model with only an intercept and no covariates. McFaddens-R^2^ provides a metric for comparison between the empirical utility model and indifference map since a numerical parameter-by-parameter comparison between ICs and iso-utility lines is not possible (random utility does not result from a parametric fit to a hyperbolic function).

### (S10) Definition and requirement criteria for revealed preference neurons

Revealed preference responses of single neurons should follow the typical scheme of behavioral ICs described by Revealed Preference Theory. These neurons comply with three simple characteristics as defined previously (Pastor-Bernier et al., 2019)

(Characteristic 1) Activity changes monotonically *across behavioral ICs* with increasing behavioral revealed preference, irrespective to bundle composition. Such monotonic neuronal response changes reflect increasing quantities of one or both bundle rewards, assuming a positive monotonic subjective value function on reward quantity.

(Characteristic 2) All bundles *along a same-revealed-preference IC* elicit the same, insignificantly varying neuronal response thus reflecting the systematic trade-off between the two bundle rewards while maintaining the same revealed preference. This crucial characteristic requires the same neuronal responses to bundles that are equally revealed preferred despite different physical bundle composition.

(Characteristic 3) The neuronal responses reflect IC non-linearities. The ICs are often not symmetric (diagonal) and linear, and the neuronal response should match these behavioral parameters. The IC slope reflects the currency relationship between the two bundle rewards, indicating the revealed preference relation between the two rewards of a bundle, and thus the value of one reward relative to a common-currency reference reward. We estimated slope and curvature parameters from regression coefficients of each neuronal response and compared them with the average slope and curvature of all behavioral ICs of a given bundle type.

We employed a combination of three statistical tests that reflected these three characteristics, comprising (1) conventional linear regressions to assess the monotonicity of neuronal response change across ICs in approximation to linearity but without formally assuming it (*P* < 0.05 for β coefficients; t-test) (see below, Eq. 4), (2) Spearman rank-correlation to confirm ordinal monotonicity of response change across ICs without assuming a particular numeric scale (*P* < 0.05), and (3) two-factor Anova to assess significant response change in the spirit of ICs: significance across ICs and insignificance within ICs, without regard of monotonicity of change (*P* < 0.05 for factors).

#### Requirement tests

Characteristic 1: All neurons with significant positive or negative changes identified by Eq. 3 needed to be also significant in the Spearman rank-correlation test.

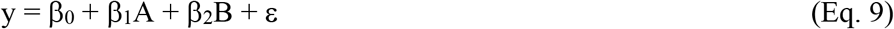

Characteristics 1 and 2: To assess together the first two necessary conditions for revealed preference coding in a direct and intuitive way, we used a two-factor Anova on each Wilcoxon-identified task-related response that was significant for both regressors in Eq. 4; the factors were across-IC (ascending rank order of behavioral ICs) and within-IC (same rank order of behavioral IC). To be a candidate for coding revealed preference, changes across-ICs should be significant (*P* < 0.05), changes within-IC should be insignificant, and their interaction should be insignificant.

Characteristic 3: Whereas the regression defined by Eq. 4 provides a conservative estimate of revealed preference coding, the full construction of neuronal ICs for comparison with behavioral ICs requires inclusion of the IC curvature that depends on both rewards. To this end, we extended Eq. 3 by adding the interaction term AB:

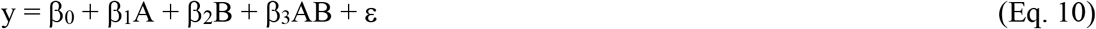

with ϵ = err_0_ + err_1_+ err_2_ + err_3_. The neuronal IC slope was estimated from the ratio of coefficients β_2_ / β_1_. Note the different meanings of the slope term: the neuronal IC slope (β_2_ / β_1_) describes the relative coding strength of the two bundle rewards (reflecting the currency of the two rewards), whereas the neuronal coding slope alone (β) describes the strength of neuronal response. The neuronal IC curvature was the β_3_ coefficient of the interaction term AB (all β’s *P* < 0.05; t-test). The regression defined by Eq. 5 is identical to the hyperbolic model used for fitting behavioral ICs (d = ax+by+cxy; Eq 1).

### (S11) Z-scoring of neuronal responses and regression against behavioural utility

To obtain a normalized count for each response, we calculated the z-score by subtracting from mean neuronal activity during the Pretrial epoch and dividing by standard deviation of that activity. We consider here a response for each unique bundle choice option corresponding to a different behavioural utility. All z-scored neuron responses (whole-window averages) in four epochs (Target onset, go, choice and reward) were independently regressed against the behavioural utility (U_b_) for each bundle

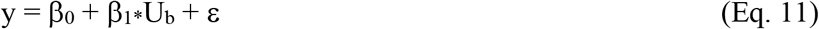

A neuron has a different β_1_ coefficient for each given task epoch, bundle and subject. Neurons that had a significant β_1_ coefficient in any epoch were investigated further. The β_1_ values and associated p-Values are described in table S1-3. Neuronal responses were obtained from the bundle blackcurrant vs grape and the bundle blackcurrant vs water in Subject 1, and bundle blackcurrant vs mango in Subject 2.

### (S12) Neuronal random utility estimation

In eq 4, Vij represents the deterministic portion of the utility: Vij = α + β*A + γ*B where α is an alternative specific constant, β is the marginal utility of component A (Blackcurrant), and γ is the marginal utility of component B (grape, grape-IMP, strawberry, water, mango, apple, yoghurt or saline). RUM uses choice data to infer β and γ. However, changes in activation in the OFC might correspond to changes in utility, and therefore they may provide a more direct measure of the marginal utilities. One way to investigate this is to incorporate the neuron response to the aggregate data in the model, where different neurons might reflect different preferences. Neurons with larger responses to a given bundle component might reflect a larger marginal utility for this component. In practice, we incorporate the responses, and the id of each individual neuron as covariates in an extended RUM multinomial model. We obtain McFadden Likelihoods to allow direct comparison between the conventional and the extended RUM model incorporating OFC responses.

### (S13) Probability and relative utility coding

In order to investigate the relationship between choice probability and neuron response, we obtained the choice probability (P) for the reference bundle option (U) and the alternative variable bundle option (W) within each binary choice block. Then, we regressed P against the z-scored responses (Y) across choice bundles against non-null bundle alternatives in two different situations:

Chosen Value W: The variable option (W) is presented for a given fixed reference option (U) in each block. W is chosen

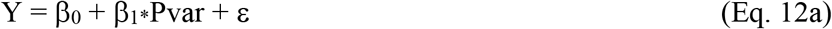

Chosen Value U: The variable options (W) is presented for a given fixed reference option (U) in each block. U is chosen

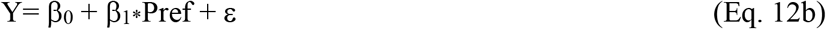

Statistically significant β regressors in Pvar and Pref identify several forms of chosen value (Table S6). Relative chosen value neuron candidates were further regressed against behavioural utility with Eq.11. Neuron RUM was computed as described in S12

### (S14) Surgical procedures and electrophysiology

A head-restraining device and a recording chamber (40 x 40mm, Gray Matter) were implanted on the skull under full general anaesthesia and aseptic conditions. The stereotactic coordinates of the chamber enabled neuronal recordings of the orbitofrontal cortex (OFC), (Paxinos et al., 2000). We located the OFC from bone marks on coronal and sagittal radiographs taken with a guide cannula inserted at a known coordinate in reference to the implanted chamber, using a medio-lateral vertical and a 20° degree forward directed approach aiming for area 13. Monkey A provided data from the left hemisphere, Monkey B from the right hemisphere, via a craniotomy in each animal ranging from Anterior 30 to 38, and Lateral 0 to 19. We conducted single-neuron electrophysiological recordings using both custom-made glass-coated tungsten electrodes (Merrill & Ainsworth, 1972), and commercial electrodes (Alpha Omega, Israel) (impedance of about 1MOhm at 1KHz). Electrodes were inserted into the cortex with a multi-electrode drive (NaN drive, Israel) with the same angled approach as used for the radiography. Neuronal signals were collected at 20kHz, amplified using conventional differential amplifiers (CED 1902 Cambridge Electronics Design) and band-passed filtered (high: 300 Hz, low: 5 kHz). We used a Schmitt-trigger to digitize the analog neuronal signal online into a computer-compatible TTL signal. However, we did not use the Schmitt-trigger to separate simultaneous recordings from multiple neurons, in which case we searched for another single-neuron recording or occasionally stored the data in analog form for off-line separation by dedicated software (Plexon offline sorter). An infrared eye tracking system monitored eye position (ETL200; ISCAN).

### (S15)Data availability

The data that support the findings of this study are available from the corresponding author upon reasonable request.

## Supplemental Info

**Figure. S1.**
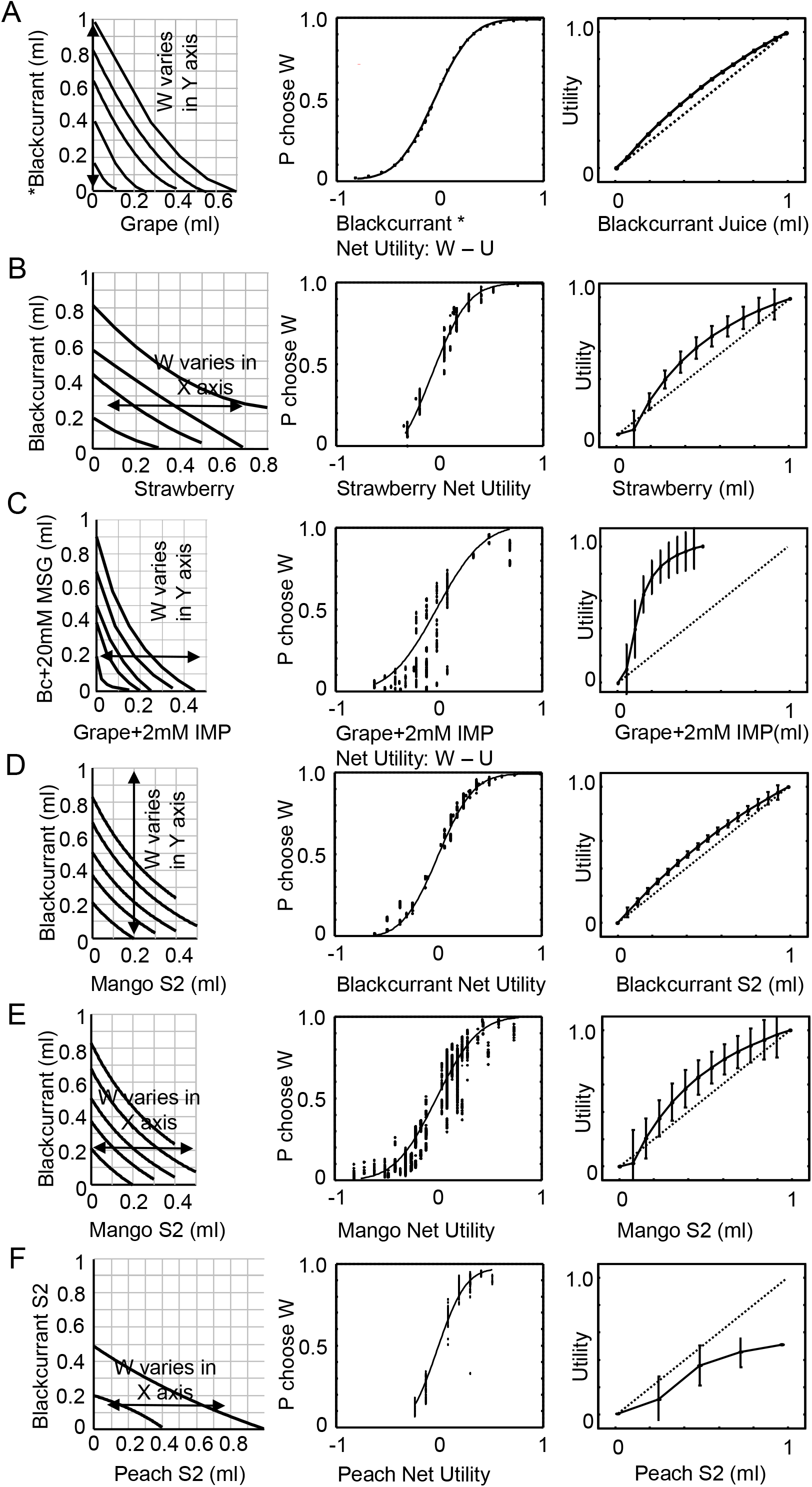
Random utility modeling on behavioral data for additional bundles in two subjects. (A) Blackcurrant vs grape control, (B) blackcurrant vs strawberry, (C) MSG-blackcurrant, vs IMP-grape,(D-E) Mango and (F) peach. The first panel on each row shows the indifference map and the direction of sampling for inference of utility. Blocks of choice trials with bundles with no grape and variable amounts of blackcurrant were used to infer the control utility function of Blackcurrant. The middle panel shows choice probability as a function of the net utility in reference and variable bundles. The right panel shows the utility function with respect to the physical amount of blackcurrant ml).The diagonal dotted line in the right panel serves to show deviation from linear utility defining risk attitude as risk averse (concave) or risk seeking (convex). B-F shows the sampling procedure, probability of choice and utility function for strawberry (B). Grape IMP (C) in Subject 1 and Blackcurrant (D), Mango (E) and Peach (F) in Subject 2.

**Figure S2.**
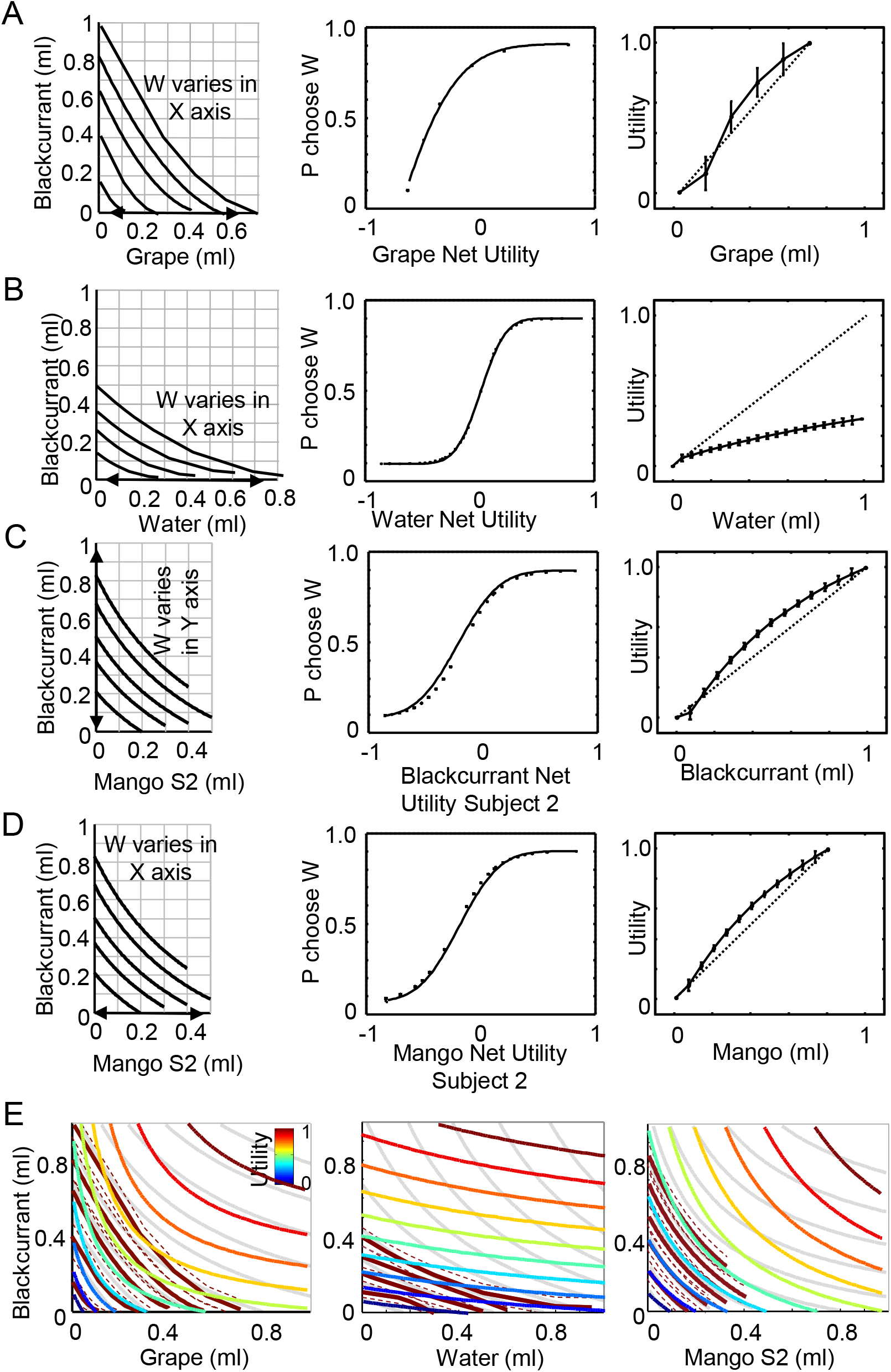
Random utility controls without interaction. Random utility modeling on behavioral data for additional bundles in two subjects: (A) Blackcurrant vs grape control, (B) blackcurrant vs water, (C) blackcurrant, vs mango,(D-E).The first panel on each row shows the indifference map and the direction of sampling for inference of utility. Blocks of choice trials with bundles with no blackcurrant and variable amounts of grape were used to infer the control utility function of Blackcurrant. The middle panel shows choice probability as a function of the net utility in reference and variable bundles. The right panel shows the utility function with respect to the physical amount of grape (ml).The diagonal dotted line in the right panel serves to show deviation from linear utility defining risk attitude as risk averse (concave) or risk seeking (convex). (E) shows comparatively the utility functions across two monkeys and three bundles, blackcurrant vs grape, blackcurrant vs water and blackcurrant vs mango. Utility maps along with empirically obtained indifference maps (black) and mixed interaction utility maps (grey) are shown for comparison.

**Figure S3.**
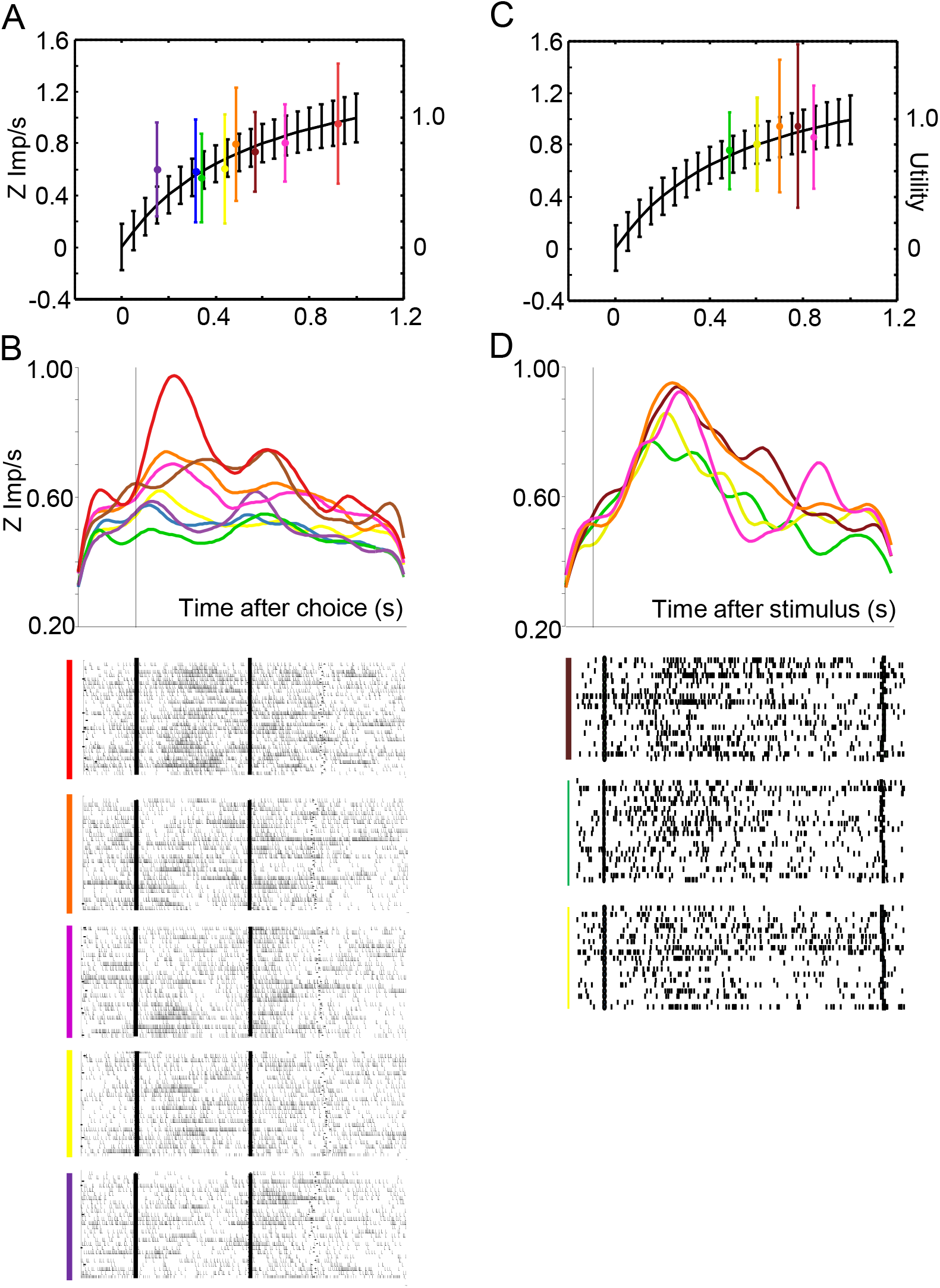
Neuronal coding of utility. (A) Z-scored responses (left axis) of an example neuron to different two-component options at Choice. The responses and SEMs are shown in different colors (red to blue color code, ranging from higher to lower). The random utility of the bundles is shown on the right axis. (B) show response histograms and representative rasters for the same neuron. (C-D) show the neuronal utility and response histograms for an additional neuron with alignment at target onset.

**Figure S4.**
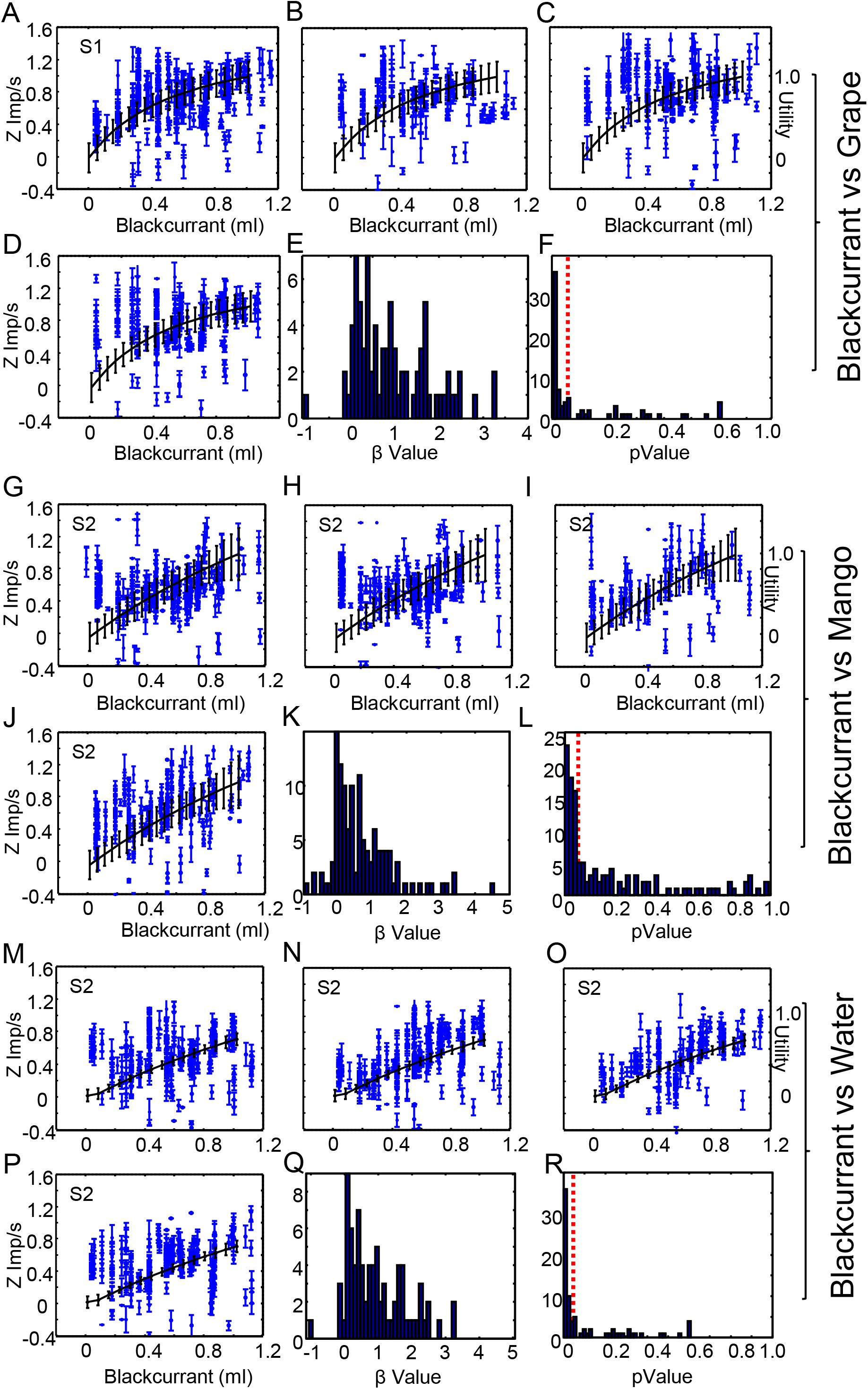
Neuronal utility in two bundles across four task epochs. (A-D) Random utility in the bundle blackcurrant vs grape in S1 at Target onset (A), GO (B), Choice (C) and Reward (D) compared to the behavioural utility function and random utility error (black).The Beta (E) and pValue (F) distributions for the responses in blackcurrant vs grape across all epochs are shown along with significance threshold (red line, p<0.05). (G-L) Random utility in the bundle blackcurrant vs mango with the same convention as above. (N-R) Random utility in the bundle blackcurrant vs water with the same convention as above.

**Figure. S5.**
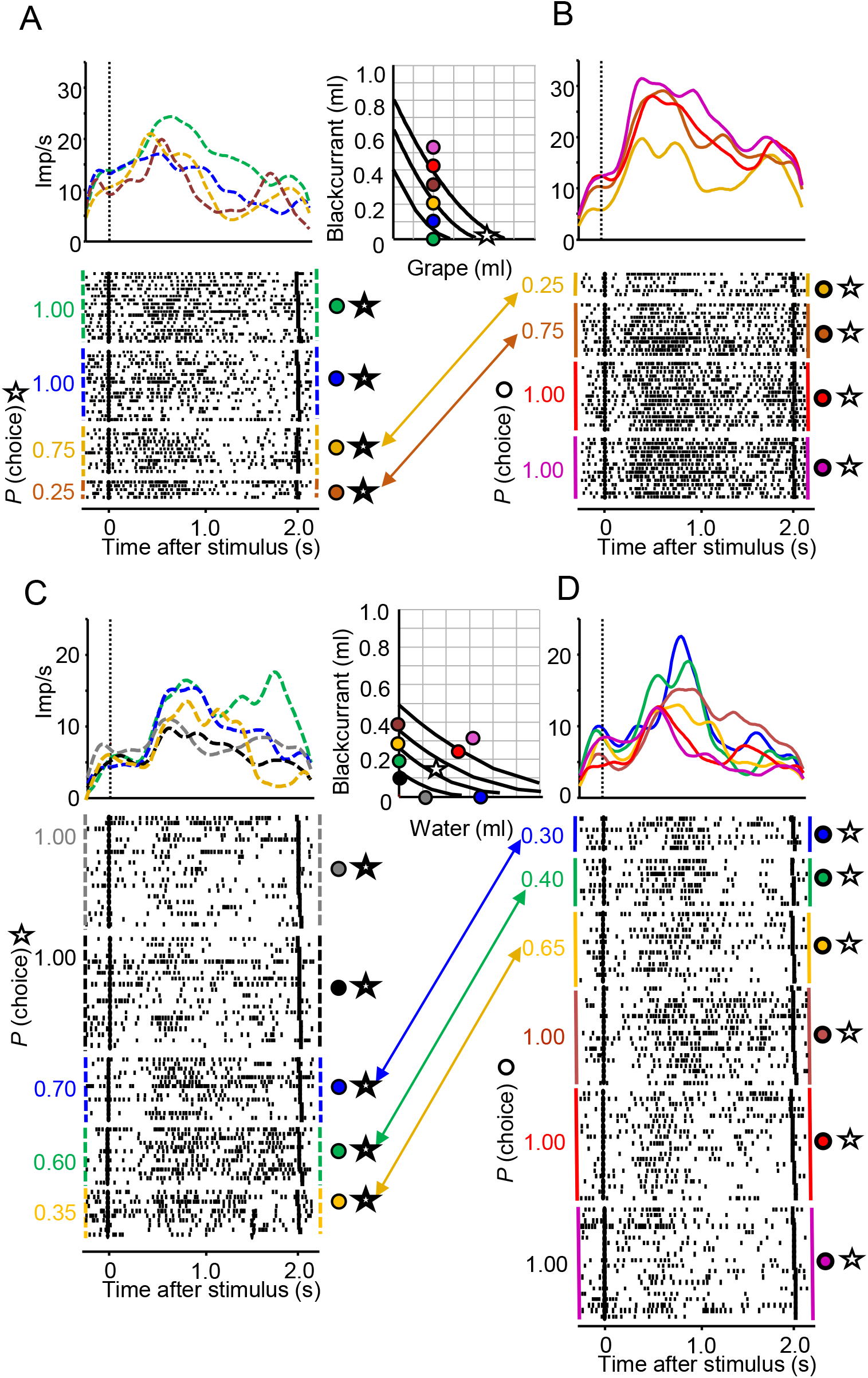
Supplementary example of relative chosen value coding and inverse relative chosen value coding in ofc. (A) Positive coding neurone showing a decrease of response for the same reference bundle chosen (star) irrespective to unchosen alternative increasing value (colored dots). (B) The same neurone shows an increase in response when the alternative bundles are chosen. Conversely, a negative coding relative chosen-value neurone (C) increases response for the same chosen reference bundle chosen (star) against an increasing unchosen value. Bundle The same neuron shows a decrease in response when the alternative bundle is chosen in (D).

**Figure S6.**
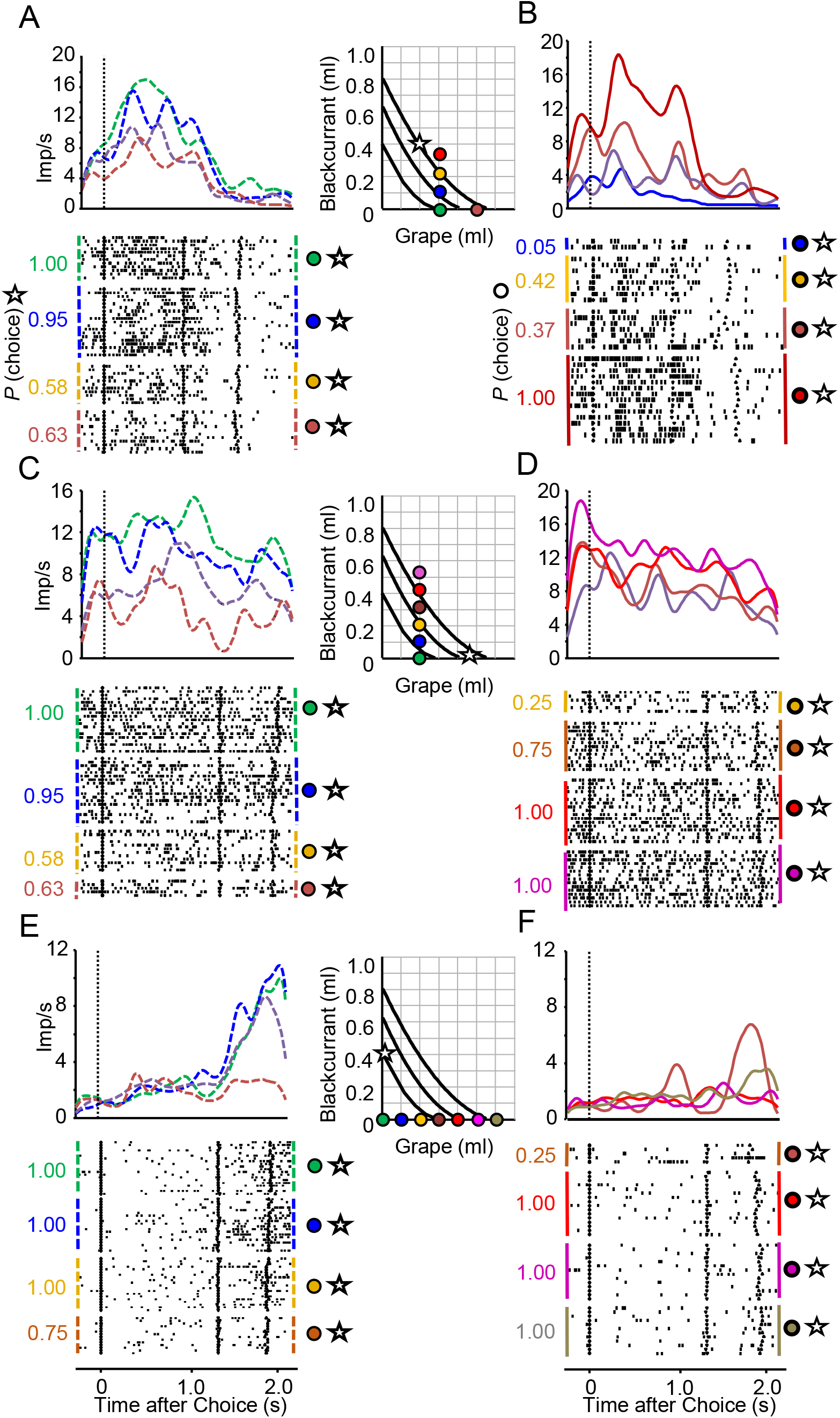
Three neurone examples of positive coding relative chosen value A. Positive coding neurones in (A, C) and (E) show a decrease of response for the same reference bundle chosen (star) against an alternative bundle of increasing value(coloured dots). The same positive coding neurons show an increase in response when the alternative bundle is chosen in (B, D) and (F). The alignment epoch corresponds to choice and reward.

## Notes

### Competing Interest Statement

The authors have declared no competing interest.

### Summary of Updates

Figure 1 has been revised along with legend and reference formatting in the main text

